# *Cis*-regulatory divergence underpins the evolution of C_3_-C_4_ intermediate photosynthesis in *Moricandia*

**DOI:** 10.1101/2021.05.10.443365

**Authors:** Meng-Ying Lin, Urte Schlüter, Benjamin Stich, Andreas P.M. Weber

## Abstract

Altered transcript abundances and cell specific gene expression patterns that are caused by regulatory divergence play an important role in the evolution of C_4_ photosynthesis. How these altered gene expression patterns are achieved and whether they are driven by *cis*- or *trans*-regulatory changes is mostly unknown. To address this question, we investigated the regulatory divergence between C_3_ and C_3_-C_4_ intermediates, using allele specific gene expression (ASE) analyses of *Moricandia arvensis* (C_3_-C_4_), *M. moricandioides* (C_3_) and their interspecific F_1_ hybrids. ASE analysis on SNP-level showed similar relative proportions of regulatory effects among hybrids: 36% and 6% of SNPs were controlled by *cis-*only and *trans-*only changes, respectively. GO terms associated with metabolic processes and the positioning of chloroplast in cells were abundant in transcripts with *cis*-SNPs shared by all studied hybrids. Transcripts with *cis*-specificity expressed bias toward the allele from the C_3_-C_4_ intermediate genotype. Additionally, ASE evaluated on transcript-level indicated that ∼27% of transcripts show signals of ASE in *Moricandia* hybrids. Promoter-GUS assays on selected genes revealed altered spatial gene expression patterns, which likely result from regulatory divergence in their promoter regions. Assessing ASE in *Moricandia* interspecific hybrids contributes to the understanding of early evolutionary steps towards C_4_ photosynthesis and highlights the impact and importance of altered transcriptional regulations in this process.

## Introduction

The majority of organic carbon in the biosphere is fixed by ribulose-1,5-bisphosphate carboxylase-oxygenase (Rubisco) through carboxylation of ribulose-1,5-bisphosphate (RuBP) during photosynthesis. However, Rubisco has affinity not only to CO_2_ but also to O_2_. The oxidation of RuBP by Rubisco generates a toxic intermediate, 2-phosphoglycolate. This by-product is metabolized through the photorespiratory pathway, which is energy-consuming and leads to the release of CO_2_. In C_4_ plants, the oxygenation reaction is repressed by the evolution of an efficient biochemical CO_2_ pump, usually functioning in concentric layers of cells known as Kranz anatomy. The enlarged bundle sheath (BS) cells with abundant organelles are located adjacent to vascular bundles and are surrounded by mesophyll (M) cells (Hatch, 1987). The CO_2_ is fixed through phospho*enol*pyruvate carboxylase in M cells and the generated C_4_ acid is decarboxylated in BS cells, where the released CO_2_ increases the CO_2_:O_2_ ratio in close proximity to Rubisco, resulting in a high carboxylation rate and a low oxygenation rate (Bräutigam & Gowik, 2016; Hatch, 1987; Schlüter & Weber, 2020). The CO_2_ compensation point is defined as the CO_2_ concentration where photosynthetic CO_2_ uptake equals CO_2_ release, *i*.*e*., no net gas exchange is detectable. C_4_ plants have much lower CO_2_ compensation points relative to C_3_ plants (Krenzer et al., 1975).

The current model of C_4_ evolution holds that C_4_ plant species evolved from the ancestral C_3_ state and C_3_-C_4_ intermediate species are considered as naturally occurring intermediates on the evolutionary trajectory towards C_4_ photosynthesis (Blätke & Bräutigam, 2019; Mallmann et al., 2014; Sage et al., 2012). C_3_-C_4_ intermediates have been reported in 21 plant lineages including dicotyledonous as well as monocotyledonous species, such as *Diplotaxis, Flaveria, Moricandia, Neurachne*, and *Panicum* (Sage et al., 2011). C_3_-C_4_ intermediate species possess a photorespiratory CO_2_ pump functioning in Kranz-like leaf anatomy, including BS cells with centripetally localized mitochondria and chloroplasts, and their CO_2_ compensation points are in between the values of C_3_ and C_4_ plants (Brown & Hattersley, 1989; Holaday & Chollet, 1984; Sage et al., 2014). The photorespiratory CO_2_ pump in C_3_-C_4_ plants efficiently recycles photorespiratory released CO_2_ via the so called glycine shuttle a.k.a. C_2_ photosynthesis (Kadereit et al., 2017; Sage et al., 2014; Schlüter & Weber, 2016). This system evolved via confining the expression of the gene encoding the P-subunit of glycine decarboxylase (GLDP) to the BS cells. Consequently, GLDP activity is absent from leaf M cells of C_3_-C_4_ plants (Monson & Rawsthorne, 2000), and hence to complete the photorespiratory pathway, glycine must be shuttled to the BS cells, where CO_2_ released from mitochondria can be efficiently recaptured by numerous, surrounding chloroplasts.

A number of anatomical and biochemical adaptive steps on the evolutionary path from C_3_ to C_4_ photosynthesis have been depicted in different models (Heckmann et al., 2013; Mallmann et al., 2014; Sage et al., 2012; Williams et al., 2013). Based on studies of various naturally occurring C_3_-C_4_ intermediate species, a stepwise model was proposed: (1) the vein density increases; (2) the leaf proto-Kranz anatomy evolves; (3) a photorespiratory CO_2_ pump is built by a reduced M:BS ratio and confinement of mitochondrial glycine decarboxylase (GDC) activity to BS cells; (4) enzymes of the C_4_ metabolic cycle are established with spatial or temporal expression adjustments of C_3_ genes (Sage et al., 2012). The consensus trajectories of the statistical (Williams et al., 2013) and mechanistic (Heckmann et al., 2013; Mallmann et al., 2014) models confirmed these steps, but the order of steps was flexible and the path was smooth (Heckmann, 2016; Williams et al., 2013). All C_4_ evolution models unequivocally predicted that the photorespiratory CO_2_ pump, resulting from the confinement of GDC activity to BS cells, is a crucial step.

In the genera *Flaveria* (*Asteraceae*) and *Moricandia* (*Brassicaceae*), the molecular mechanisms by which *GLDP* expression becomes confined to BS cells during evolution of C_3_-C_4_ intermediacy have been resolved. The genomes of C_3_ *Flaveria* species encode two isoforms of *GLDP*, one BS-specific isoform (*GLDPA*) and the other one ubiquitously expressed in all photosynthetic tissues (*GLDPB*). *GLDPB* becomes a pseudogene in C_3_-C_4_ intermediate *Flaverias* and thereby GDC activity is lost from M cells during C_4_ photosynthesis evolution (Schulze et al., 2013). A conceptually similar mechanism underpins the independent but convergent evolution of C_3_-C_4_ intermediacy in the *Brassicaceae*. The promoter of the *GLDP1* gene in this plant family carries two conserved *cis*-regulatory elements, one that drives expression in the M cells (M-box), and another one that governs expression in the vasculature (V-box). Deletion of the M-box from the *GLDP1* promoter led to the restriction of GDC activity to BS cells (Adwy et al., 2015; Adwy, 2018). The establishment of the C_3_-C_4_ intermediate photorespiratory CO_2_ pump very likely requires further metabolic adjustments and anatomical modifications, probably through altered transcriptional regulation. However, the genetic factors underpinning this process are still mostly unknown.

*Cis*-regulatory divergence has been reported to play an important role in adaptive phenotypic evolution because, compared to the nonsynonymous mutation in protein sequences, it causes fewer deleterious pleiotropic effects (Stern & Orgogozo, 2008; Wittkopp & Kalay, 2012; Wray, 2007). Additionally, *cis*-regulatory divergences have impacts on limiting gene expression to particular tissue or cellular compartments, to specific life stages or environments (Prud’homme et al., 2007). Allele specific expression (ASE) analysis on heterozygote sites in diploid hybrids is considered as an effective method to identify *cis-*acting factors, as allelic expressions are under the same feedback control and sharing non-*cis*-elements. Comparing the allelic ratio between parental alleles and that in hybrids could distinguish the effect between *cis*- and *trans*-factors (Li et al., 2017). Progress in sequencing technologies, next-generation sequencing (NGS)-based approaches, such as RNA-Seq, enables analyzing ASE on a transcriptome-wide scale. This strategy has been widely applied to yeast, fruit flies, and also plants, including *Arabidopsis, Capsella, Atriplex*, maize, rice, millet (He et al., 2012; Lemmon et al., 2014; McManus et al., 2010; Rhoné et al., 2017; Shao et al., 2019; Steige et al., 2015; Sultmanis, 2018; Tirosh et al., 2009). Here we apply ASE analysis on interspecific hybrids of parents displaying C_3_ and C_3_-C_4_ intermediate photosynthesis.

The genus *Moricandia* provides an outstanding system to unravel the regulatory mechanisms underpinning C_3_-C_4_ intermediacy. It comprises species with C_3_ and C_3_-C_4_ photosynthesis existing in a close phylogenetic proximity. *Moricandia* species share phylogenetic proximity with species in other Brassicaceae genera, such as *Brassica* and *Diplotaxis*, as well as with Arabidopsis, and were close to C_4_ species in the family *Cleomaceae* (Kellogg, 1999; Beilstein et al., 2010; Schlüter et al., 2017). To study the inheritance of *Moricandia* C_3_-C_4_ characteristics, intergenic hybridizations of *Moricandia* with distant *Brassica* relatives have been reported in the literature, through embryo rescue, sexual crosses and somatic hybridizations (list in Warwick et al., 2009). Interspecific hybrids of *Moricandia* C_3_ and C_3_-C_4_ species were generated by *M. arvensis* as maternal and *M. moricandioides* as paternal species, of which the phenotypic characterization is so far limited to records of CO_2_ compensation points (Apel et al., 1984). Most experimental hybrid studies were abandoned at that time, because of reproductive disorders. However, genomic studies are facilitated now by the access to genetic resources of closely related *M. moricandioides* (C_3_) and *M. arvensis* (C_3_-C_4_) (Lin et al., 2021). In this study, we assessed ASE on SNP- and transcript-level in *Moricandia* by means of RNA-Seq on *M. arvensis* (C_3_-C_4_), *M. moricandioides* (C_3_), and six of their interspecific F_1_ hybrids. Gene ontology assessments identified genes participating in chloroplast relocation and genes demonstrating extreme allele imbalance. The spatial gene expression pattern of selected genes flagged by ASE was validated by promoter-GUS analysis in *A. thaliana*. Our results indicate a predominance of *cis*-regulatory effects on early evolutionary steps of C_4_ photosynthesis.

## Results

### *Moricandia* interspecific hybrids display phenotypes between that of parental genotypes

For the analysis presented here, interspecific hybridization in *Moricandia* was performed using *M. arvensis* (C_3_-C_4_) as maternal and *M. moricandioides* (C_3_) as paternal species (Ma×Mm). The reciprocal cross, Mm×Ma, had produced only one fifth of the seeds compared to Ma×Mm hybridization. Additionally, the germination rate of seeds from Ma×Mm and Mm×Ma was 86% and 25%, respectively. In *M. arvensis* leaves, organelles are found in the BS and M cells along the inner tangential walls and are abundantly accumulated toward veins in the BS cells (Beebe & Evert, 1990; Schlüter et al., 2017). The same leaf anatomy was observed in this study: chloroplasts are not only arranged on the inner wall of M cells, but also abundantly accumulated towards veins in BS cells in *M. arvensis*. In contrast, only few chloroplasts were found evenly distributed along the inner wall in BS cells, and some on the inner wall of M cells in *M. moricandioides* (Figure 1). The CO_2_ compensation points of *M. arvensis* and *M. moricandioides* were 23.6±2.7 and 55.6±3.2 ppm, respectively, consistent with previous studies (Apel, 1980; Bauwe & Apel, 1979; Schlüter et al., 2017). The *Moricandia* interspecific hybrids displayed variation in their CO_2_ compensation points, ranging from 39 to 55 ppm, generally between the parental lines, but closer to that of C_3_ species (Figure 2). These hybrids further varied for the amount and arrangement of chloroplasts in BS cell: chloroplasts were evenly distributed along the inner wall in BS cells of hybrid 1, 2, 5, 6, but only few chloroplasts were found in BS cells of hybrid 3, 4 (Table 1, Supplementary file 1). The non-uniformity of hybrids regarding the various phenotypic characteristics might be due to the heterozygosity and chromosome mismatching of their parental species. Increased vein density compared to C_3_ species was found in *Heliotropium* and *Flaveria* C_3_-C_4_ species (Muhaidat et al., 2011; Sage et al., 2013), but not in *Moricandia* (Schlüter et al., 2017). The leaf venation was observed from the top view of cleared leaves under the light microscope and the vein density was calculated as the vein length per area. The vein density of *M. arvensis* was not significantly higher than that of *M. moricandioides* but broader veins were observed in *M. arvensis* because of more chloroplasts accumulating toward vascular bundles in BS cells (Supplementary file 1 and 2). The vein density of hybrids showed no differences compared to parental species and the leaf venation of hybrids was more similar to that of the C_3_ parent with thinner veins corresponding to their leaf anatomy (Supplementary file 2). They produced only very few F_2_ seeds, likely because of abnormal pollen produced by the F_1_s, resulting in sterility of hybrids (Supplementary file 3). Many interspecific or intergeneric hybrids were reported to be sterile as a result of abnormal chromosome pairing or irregular meiotic division of pollen mother cells (Apel et al., 1984; H. R. Brown & Bouton, 1993; Covshoff et al., 2014). Taken together, the *Moricandia* interspecific hybrids demonstrated intermediate characteristics of CO_2_ compensation points and leaf anatomy between that of C_3_ and C_3_-C_4_ parents; however, they more resembled the C_3_ species. The vein width of hybrids was similar to that of C_3_ plants.

**Figure 1.**
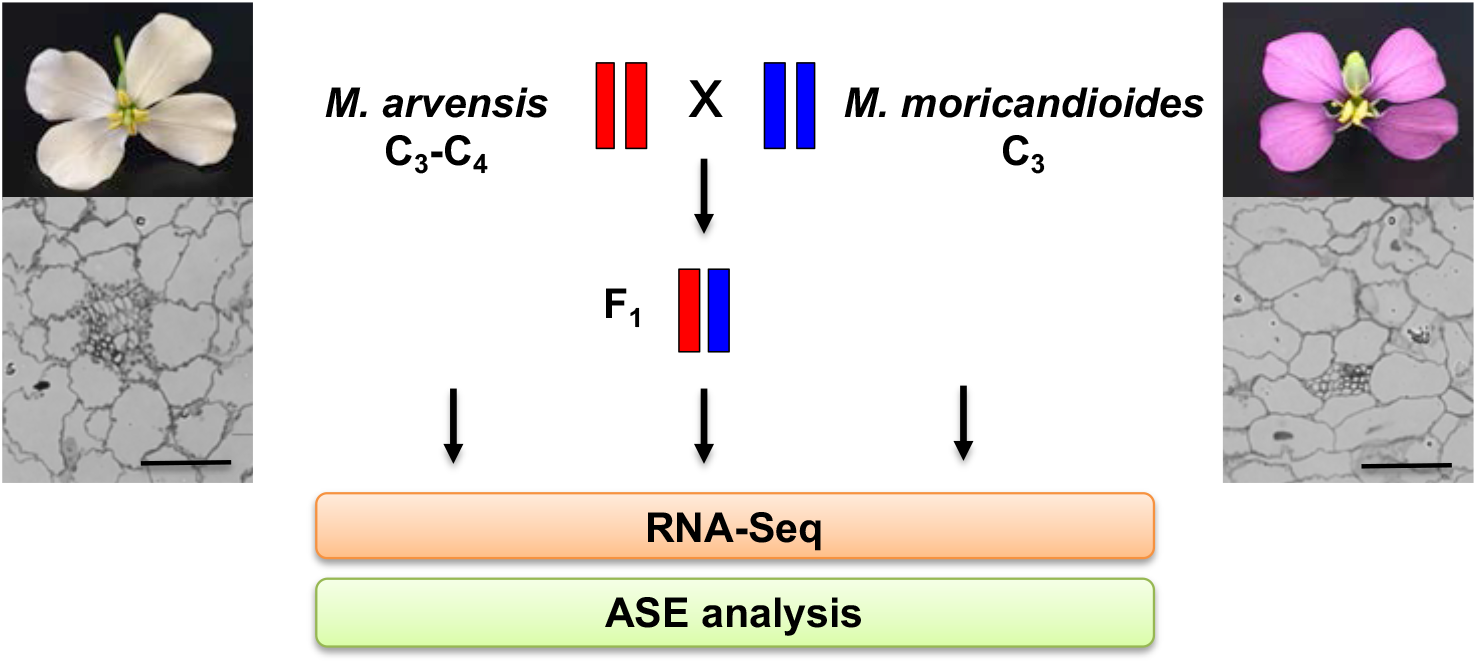
Experimental design. The interspecific hybrids were obtained from the hybridization of *M. arvensis* as maternal and *M. moricandioides* as paternal species. Leaf cross sections are shown below the pictures of typical flowers of the parental lines. RNA-Seq was processed on parents and selected hybrid lines, and further introduced to ASE analysis. Bar, 100 µm.

**Figure 2.**
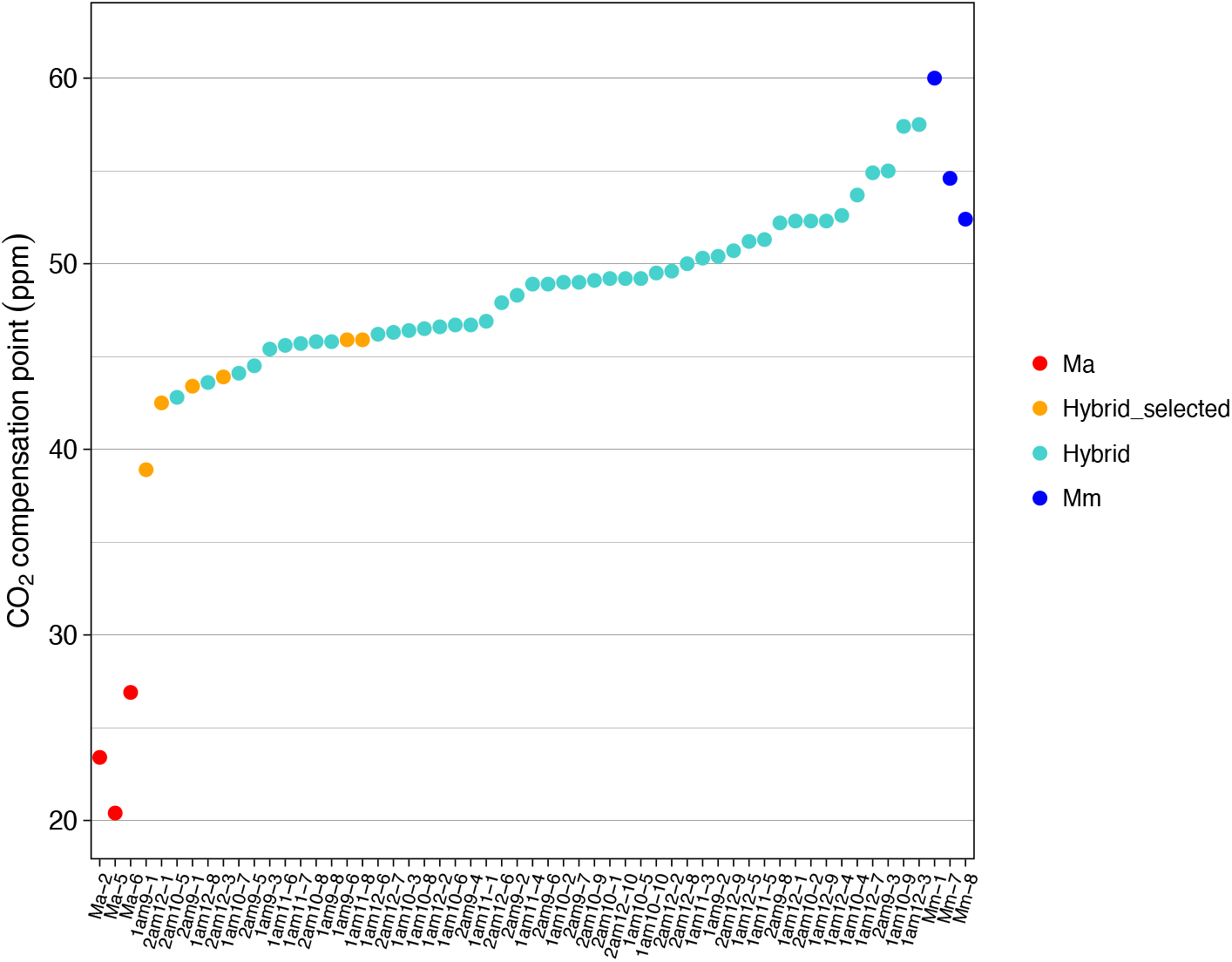
Distribution of CO_2_ compensation points of *M. arvensis, M. moricandioides* and their interspecific hybrids. See ***Figure 2—source data 1*** for values of CO_2_ compensation points. Ma, *M. arvensis*; Mm, *M. moricandioides*; Hybrid, in total 51 hybrids were tested for their CO_2_ compensation point; Hybrid_selected, six interspecific hybrid lines with relative low value processed to RNA-Seq analysis. **Source data 1**. CO_2_ compensation points of *M. arvensis, M. moricandioides* and their interspecific hybrids.

**Table 1.** Phenotypic characterization of *Moricandia* parental species and their interspecific hybrids applied to RNA-Seq analysis. *M. arvensis* and *M. moricandioides* demonstrated typical C_3_-C_4_ and C_3_ phenotypes, respectively. Interspecific hybrid lines indicated not uniform characteristics, generally intermediate between that of parents, but more resemble to C_3_ parent.

| Sample | Sample name | CO<sub>2</sub> compensation point (ppm) | Chloroplasts in BS cells |
| --- | --- | --- | --- |
| *M. arvensis* | Ma | 24.99 | Abundant, centripetally accumulated |
| *M. moricandioides* | Mm | 54.91 | Few, Centripetally and centrifugally accumulated |
| I am 9-1 | Hybrid1 | 38.92 | Centripetally and centrifugally accumulated |
| II am 12-1 | Hybrid2 | 42.53 | Centripetally and centrifugally accumulated |
| II am 9-1 | Hybrid3 | 43.38 | Nearly none |
| II am 12-3 | Hybrid4 | 43.89 | Nearly none |
| I am 9-6 | Hybrid5 | 45.87 | Centripetally and centrifugally accumulated |
| I am 11-8 | Hybrid6 | 45.93 | Centripetally and centrifugally accumulated |

### Transcript patterns showed no strong differences between C_3_ and C_3_-C_4_ *Moricandia*, but transcripts of genes predicted to be involved in the glycine shuttle tended to be enhanced in the C_3_-C_4_ species

The transcriptome of *M. arvensis* (C_3_-C_4_) and *M. moricandioides* (C_3_) was assembled using STAR v.2.5.2b with the draft genome of *M. moricandioides* serving as the reference (Lin et al., 2021). Three replicates from *M. arvensis* showed 66% mapping rate, and three replicates from *M. moricandioides* showed 94% mapping rate. Mapping rates of hybrids on *M. moricandioides* ranged from 61 to 81%. Principle component analysis (PCA) showed that the first principle component (PC1) explained 72% of the variance and clearly separated samples by species (Supplementary file 4A). PC2 underlined the separation of three replicates of *M. moricandioides* (Supplementary file 4A). The assessment of differential gene expression on 35,034 transcripts was performed with the DESeq2 tool (Love et al., 2014). Transcripts with a false discovery rate (FDR) ≤ 0.01, *P*-value adjusted with the BH procedure, were annotated as significantly differentially expressed. Using this definition, we found 3,491 transcripts that were significantly differentially expressed in *M. arvensis* and *M. moricandioides* leaves, where 2,712 transcripts were downregulated and 779 transcripts were upregulated in the C_3_-C_4_ species *M. arvensis*. GO terms such as metabolic process of small molecule, organic acid, and carbohydrate, transport of water and fluid, Golgi/endomembrane system organization were found in transcripts upregulated in C_3_-C_4_ *Moricandia* (Supplementary file 5). The downregulated transcripts encompassed significantly overrepresented GO terms, such as telomere maintenance, meiotic/nuclear chromosome segregation, chromosome organization, and regulation of organelle organization (Supplementary file 5).

The metabolic difference between C_3_ and C_3_-C_4_ plants is predominantly caused by different intercellular arrangement of the photorespiratory process. Therefore, genes involved in pathways, such as glycine shuttle, C_4_ cycle, Calvin-Benson cycle, and mitochondrial e^-^ transport, were screened for evidence of differential expression (Supplementary file 6). 12 out of 36 genes in the glycine shuttle were upregulated in C_3_-C_4_ *Moricandia*, such as *PGLP2, PLGG1, HPR1, SHMT2, GLDT, GLHD3, GS2*. 14 out of 59 genes in C_4_ cycle were upregulated in C_3_-C_4_ *Moricandia*, such as *alpha CA1, gamma CA2, NADP-MDH, BASS2, AspAT2, NAD-ME1, PEPCK*. Out of 40 genes in the Calvin-Benson cycle, only *GAPA2* was upregulated in C_3_-C_4_ *Moricandia*. Out of 5 genes in mitochondrial e^-^ transport, *UCP1* and *UCP2* were upregulated in C_3_-C_4_ *Moricandia*. All in all, upregulated transcripts were overrepresented in glycine shuttle, C_4_ cycles, and mitochondrial e^-^ transport relative to upregulated genes in all pathways in C_3_-C_4_ intermediate species (Chi-squared test, adjusted P-value ≤ 0.01), supporting the current models for glycine shuttle pathways in C_3_-C_4_ intermediates (Supplementary file 6).

### ASE analysis on SNP-level showed similar relative proportions of regulatory effects among hybrids

Gene expression regulation is governed by the interaction of distinct regulatory effects (*cis*- and *trans*-acting factors). In hybrids, the two alleles inherited from the parental genotypes are under the same cellular conditions and shared non-*cis*-elements. In this study, interspecific hybrids of *M. arvensis* and *M. moricandioides* were utilized for discovering transcriptional regulation through ASE analysis. Comparison of the allele ratio of characteristic SNPs between hybrids and parents enabled us to distinguish between *cis*- and *trans*-regulatory effects (Li et al., 2017). SNPs indicating *cis-*only regulatory effects (*cis*-SNP) are those for which the two alleles are expressed unequally in hybrids and the allele ratio is the same between parents and hybrids. SNPs with *trans-*only regulatory effects (*trans*-SNP) are defined by equal allele expression in hybrid, but unequal in parents. To assess ASE between *M. arvensis* and *M. moricandioides*, three replicates of each parental species and six interspecific hybrids were selected for RNA-Seq (Figure 1). The ASE analysis on six hybrids was conducted individually on a set of 147,883 SNPs in *Moricandia* (Figure 3—source data 1 and source data 2). The six interspecific hybrids displayed similar relative proportions of regulatory effects (Figure 3). On average, 36% of SNPs showed regulatory divergence by *cis-*only regulatory effect (*cis*-SNP), and 7% of SNPs indicated *trans-*only effects (*trans*-SNP). Furthermore, 21% of SNPs indicated mixed effects (*cis-* plus *trans-*SNP) and 37% of SNPs showed neither *cis-* nor *trans-*regulatory effects (no *cis*-no *trans*-SNP). *GLDP1* of *Moricandia*, known for cell specific expression between M and BS cells regulated by the M-box in the promoter region, was tagged by *cis*-SNPs in all hybrids. Although relative proportions of regulatory effects were similar among hybrids, most cases of ASE-SNPs were specific to individual interspecific hybrids. Around 8.7% of *cis*-SNPs were shared by all hybrids (3,142 transcripts harboring common *cis*-SNPs), and only 1% of *trans*-SNPs (62 transcripts harboring common *trans-*SNPs) were shared by all hybrids. Overall, *cis*-acting effects dominated the gene expression regulation in *Moricandia* interspecific hybrids.

**Figure 3.**
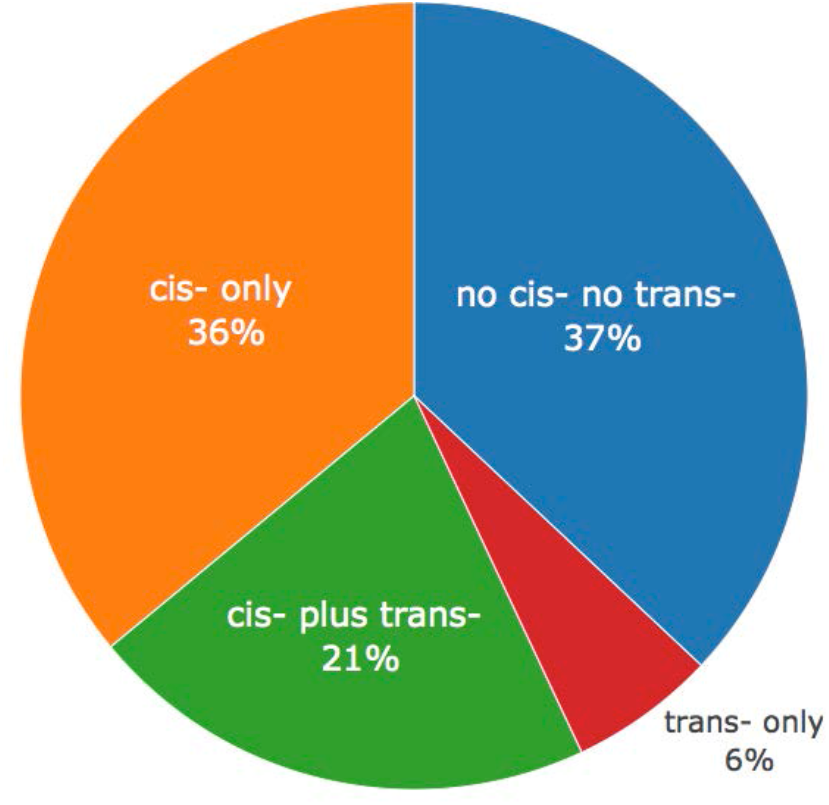
Average relative proportions of regulatory effects on SNP sites among six *Moricandia* interspecific hybrids. See ***source data 1*** for number of SNP sites defined with regulatory effects among six interspecific hybrids and ***source data 2*** for the raw data of ASE analysis based on SNP level. **Source data 1**. Regulatory effects on SNP sites among six *Moricandia* interspecific hybrids. **Source data 2**. Raw data of ASE analysis based on SNP level.

### Gene ontology (GO) and pathway enrichment analysis on transcripts with common *cis*-SNPs and common *trans*-SNPs

GO and pathway enrichment analysis were used to annotate the functions of transcripts with common *cis-*SNPs and common *trans-*SNPs. A custom-mapping file was created, containing *Moricandia* transcript names and the corresponding GO terms derived from *A. thaliana* genes. The 2,236 and 45 *Moricandia* transcripts with common *cis-*SNPs and common *trans*-SNPs, respectively (Supplementary file 7), were processed with the custom mapping file by topGO R-package for gene set enrichment analysis of biological processes (Alexa et al., 2006). The top 30 most significantly enriched GO terms for transcripts with common *cis*-SNPs relate to isopentenyl diphosphate biosynthesis, carbohydrate catabolic process, oxidoreduction coenzyme metabolic process, and chloroplast relocation (Supplementary file 8). GO terms related to nucleosome assembly, RNA methylation, organophosphate biosynthetic process, and peptide metabolic process were abundant in transcripts with common *trans*-SNPs (Supplementary file 8).

To further decipher biosynthetic pathways in which transcripts with common *cis*-SNPs and common *trans*-SNPs participate, pathway enrichment analysis was conducted using the KEGG Orthology Based Annotation System (KOBAS) (Xie et al., 2011). Transcripts with common *cis-*SNPs were significantly (P<0.05) enriched in 27 pathways, including carbon metabolism, protein processing in endoplasmic reticulum, carbon fixation in photosynthetic organisms, porphyrin and chlorophyll metabolism, glyoxylate and dicarboxylate metabolism, and nitrogen metabolism (Supplementary file 9). In contrast, transcripts with common *trans-*SNPs were significantly enriched in 9 pathways, which were related to ribosome, carbon metabolism, biosynthesis of amino acid and secondary metabolites, and fatty acid metabolism/degradation/biosynthesis (Supplementary file 9).

Therefore, the GO and pathway enrichment results suggested that *Moricandia* transcripts with *cis* mechanisms play a more prominent role in C_3_-C_4_ related functions such as major photosynthetic pathways and chloroplast relocation, whereas transcripts with *trans* mechanisms are involved in more general biological pathways.

### Transcripts with *cis*-specificity showed biased expression toward C_3_-C_4_ species in *Moricandia*

*cis*-regulatory divergences generally dominate adaptive evolution because they tend to cause fewer deleterious pleiotropic effects than nonsynonymous mutations in protein-coding sequences (Stern & Orgogozo, 2008; Wittkopp & Kalay, 2012; Wray, 2007). Further, they frequently cause altered spatiotemporal gene expression patterns (Prud’homme et al., 2007). The compartmentation of CO_2_ assimilatory enzymes between BS and M cells in C_4_ plants results from modifications in regulatory sequences (Gowik et al., 2004; Sheen, 1999). Thus, genes with *cis*-specificity (*cis*-SNPs or *cis-* plus *trans*-SNPs) were candidates for selections of direct targets or promotions of spatial gene expression during C_4_ evolutionary trajectories. In hybrid 1, we observed 4,684 *cis*-specificity SNPs (1,105 transcripts) expressed toward *M. arvensis* (Ma-biased) and 3,871 SNPs (820 transcripts) expressed toward *M. moricandioides* (Mm-biased). Similar proportions were observed in the other interspecific hybrids (Supplementary file 10). In total, there were 513 Ma-biased and 326 Mm-biased transcripts with *cis*-specificity shared by all hybrids.

To understand the gene function of biased transcripts with *cis*-specificity, the common Ma-biased and common Mm-biased transcripts were further investigated by GO term classification using Web Gene Ontology Annotation (WEGO) software (Ye et al., 2018). These transcripts were classified into GO terms under biological process, cellular component, and molecular function categories (Figure 4). There were more transcripts enriched in GO terms, such as anatomical structure morphogenesis, biosynthetic process, localization, endomembrane system, catalytic activity, and transmembrane transporter activity, in common Ma-biased than in common Mm-biased transcripts. Additionally, under anatomical structure morphogenesis, compared to common Mm-biased transcripts, more common Ma-biased transcripts were enriched in leaf morphogenesis. Therefore, this directional expression bias indicated that *cis*-specificity caused enhanced expression of C_3_-C_4_ genes involved in leaf anatomical structure organization and activity of transmembrane transporters, which could be involved in shaping C_3_ to C_3_-C_4_ photosynthesis.

**Figure 4.**
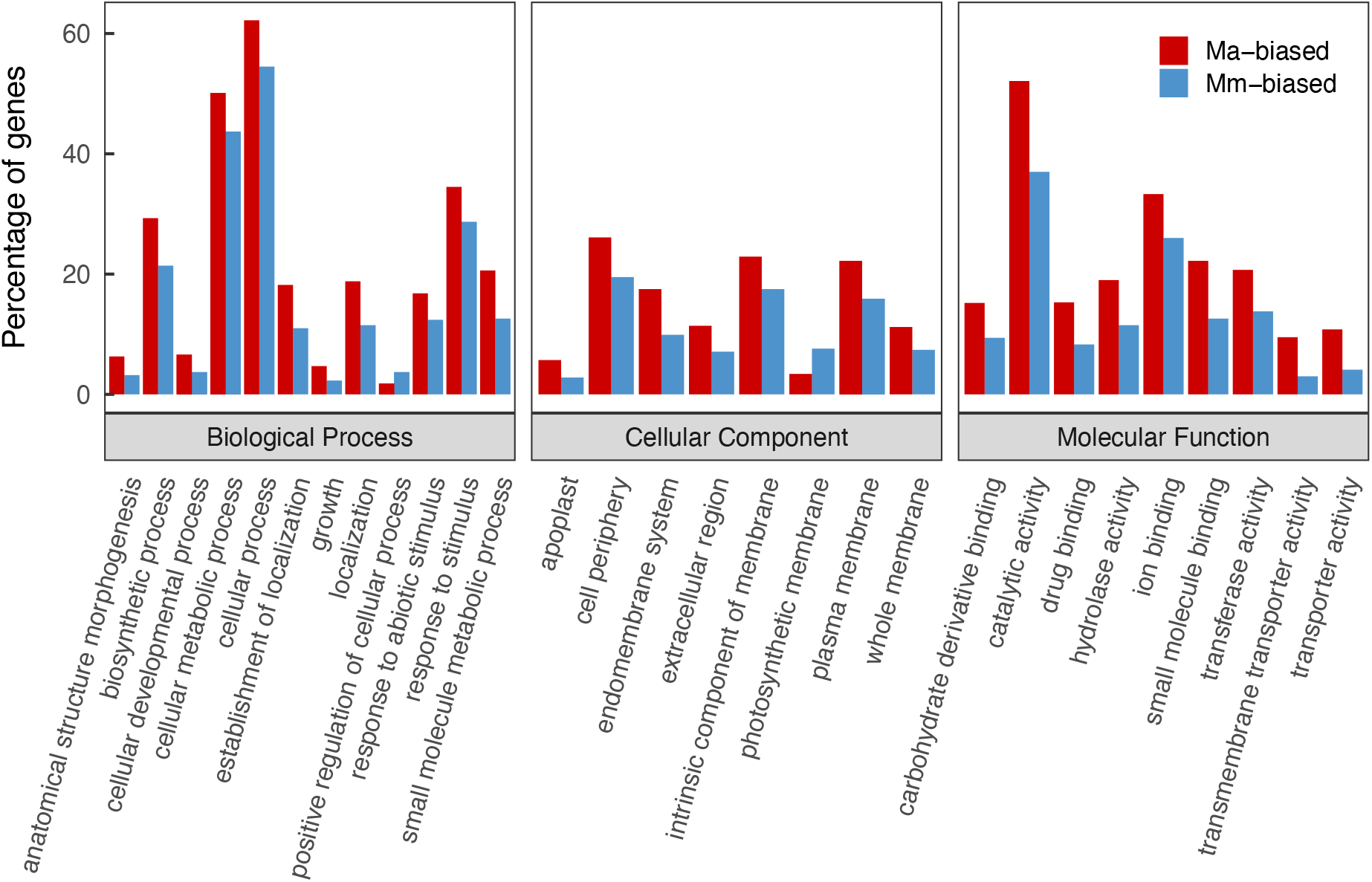
GO terms comparison between transcripts with *cis*-specificity biased toward *M. arvensis* (Ma-biased) and *M. moricandioides* (Mm-biased). All GO terms shown here were significantly different in gene percentages between Ma-biased and Mm-biased transcripts.

### Evaluating ASE on transcript-level in *Moricandia* interspecific hybrids

Assessing ASE on SNP-level allows assigning SNPs into four categories of regulatory effects. However, SNPs within a transcript might indicate different regulatory effects. For instance, *GLDP1* in hybrid 1 revealed 5 *cis*-SNPs, 16 *cis*-plus *trans*-SNPs, and 2 *trans*-SNPs. Therefore, studies evaluating ASE on SNP-level require either phased information of read counts at SNPs from genomic data of hybrids (He et al., 2012; Rhoné et al., 2017; Skelly et al., 2011; Steige et al., 2015) or an agreement across SNPs in the same transcript (Shao et al., 2019). In our study, the genomic information of hybrids was not available. Therefore, we followed the second strategy and applied a meta-analysis based allele-specific expression detection (MBASED). This method assigns the allele with higher read counts at each SNP to the major allele as a pseudo-phasing approach and therewith assumes a consistent direction of ASE within a transcript. The ASE level of transcripts was estimated with the major allele frequency (MAF), ranging from 0.5 to 1.0 (Mayba et al., 2014). The distribution of MAF was right-skewed with the mode of 0.6, implying that most of transcripts showed mild allelic imbalance (Figure 5). Transcripts with MAF ≥ 0.7 and adjusted *P*-value ≤ 0.05 were defined as ASE-transcripts. Across the six interspecific hybrids, around 27% of examined transcripts showed evidence of ASE, where 3% of them had extreme allelic imbalance (MAF ≥ 0.9 and adjusted *P*-value ≤ 0.05) (Figure 5—source data 1). *GDLP1* in *Moricandia*, known for cell specific expression regulated through *cis*-regulatory elements, showed an MAF of 0.76 on average across the six hybrids (Figure 5—source data 2). Interestingly, some transcripts showed extreme allele bias to one of the parental species in hybrids (Supplementary file 11). For instance, *MSTRG*.*16015* encoding a putative chloroplast RNA binding protein showed an average MAF of 0.96 toward *M. arvensis* allele, and *MSTRG*.*5109* encoding ATP synthase subunit d revealed an average MAF of 0.96 biased to the *M. arvensis* allele (Figure 5—source data 2). Overall, 27% of assayed transcripts were defined as ASE-transcript and a group of transcripts had strong allelic imbalance in hybrids.

**Figure 5.**
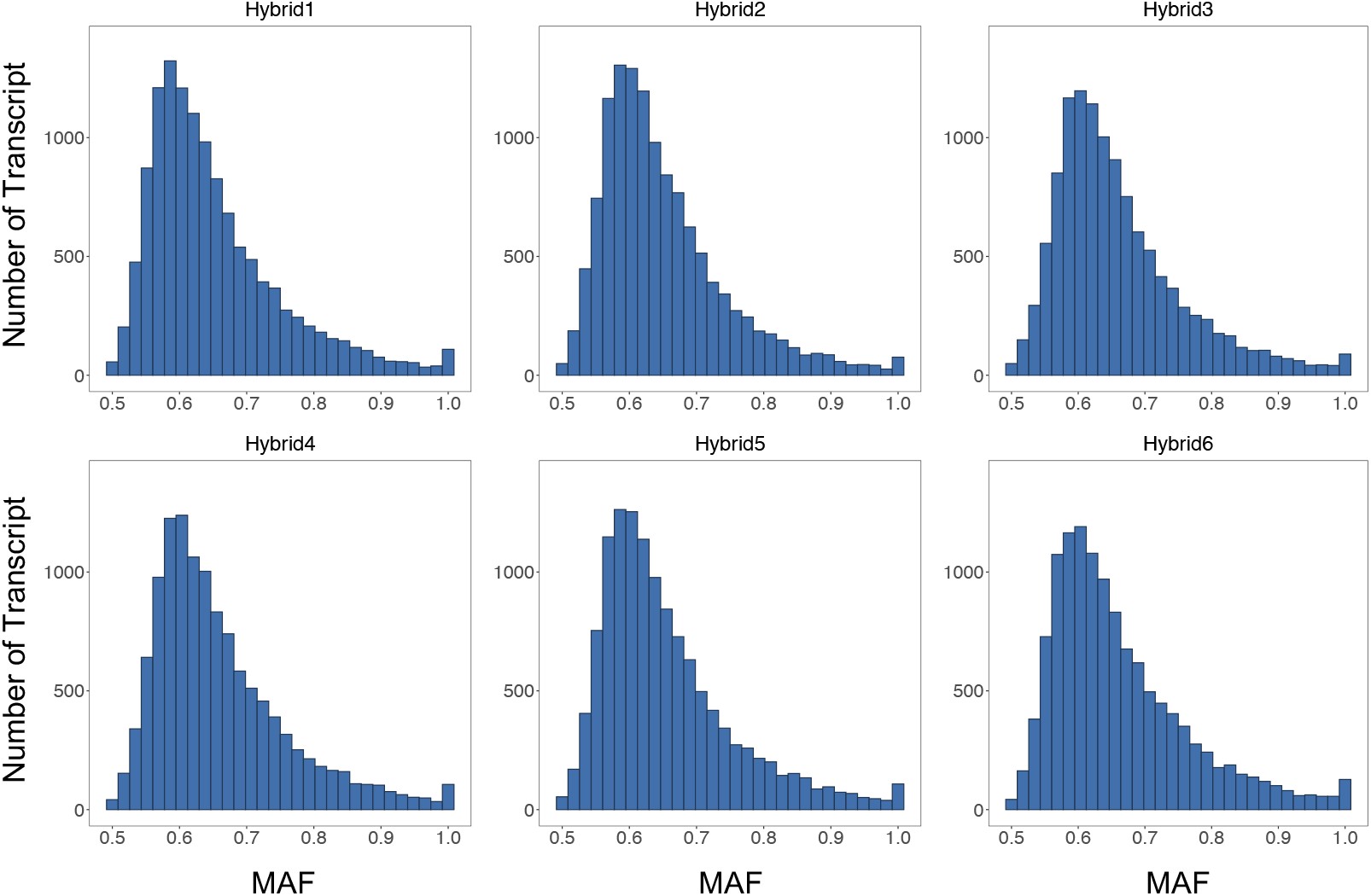
Distribution of allelic imbalance for all assayed transcripts among six *Moricandia* hybrids. The major allele frequency (MAF) represented the intensity of allelic imbalance in hybrids, obtained by MBASED (Mayba et al., 2014), using the allele with more read counts as major allele. See ***source data 1*** for number of assayed transcripts among six hybrids and ***source data 2*** for the list of transcripts with MAF among six hybrids and ***source data 3*** for the ***r***aw data of ASE analysis based on transcript level. Transcripts with MAF ≥ 0.7 and adjusted p-value ≤ 0.05 were defined as ASE-transcripts. Around 27% of transcripts was with MAF ≥ 0.7 and 3% of them showed MAF ≥ 0.9. **Source data 1**. Allelic imbalance for all assayed transcripts among six *Moricandia* hybrids. **Source data 2**. The major allele frequency of six *Moricandia* hybrids. **Source data 3**. Raw data of ASE analysis based on transcript level.

### Enrichment of regulatory divergences in selected pathways

Most of *Moricandia* transcripts indicated no significant differential expression in comparative transcriptome studies between the parental species using total leaf extracts. However, *cis*-acting factors, regulating spatiotemporal gene expression and transcriptional abundance, play a crucial role in adaptive phenotype evolution (Lemmon et al., 2014; Wray, 2007). Therefore, we examined the enrichment of regulatory effects on transcripts in selected pathways through detecting *cis*-SNPs and evaluating allelic imbalance in transcripts. Out of selected 140 genes, the expression of 105 genes in glycine shuttle, C_4_ cycle related, and Calvin-Benson cycle indicated altered *cis* regulation with the evidence of at least one *cis-*SNP or common *cis-*SNP (Supplementary file 12). In addition, the ASE evaluated on transcript-level demonstrated the intensity of allelic imbalance in hybrids. Transcripts with major alleles frequency ≥ 0.7 were considered to possess ASE. Out of 41 genes in the C_3_-C_4_ glycine shuttle, 32 genes indicated allele specific expression, such as *PLGG, GOX, GGAT, SHMT, GDC* complex, *DIT*, and *GS2* (Figure 6A). Furthermore, C_4_ cycle genes, such as *CA, PEPC, PPT, NADP-MDH, DIT, NADP-ME, BASS2, PPDK, PPT, AspAT*, and *PEPCK*, and Calvin-Benson cycle genes, except *SBPase*, indicated ASE (Figure 6B and 7). Taken together, most genes involving in selected pathways revealed no strong differential gene expression between C_3_ and C_3_-C_4_ species in *Moricandia*, however regulatory divergences play an important role in different photosynthetic types by ASE of critical genes involving in early C_4_ photosynthesis evolution.

**Figure 6.**
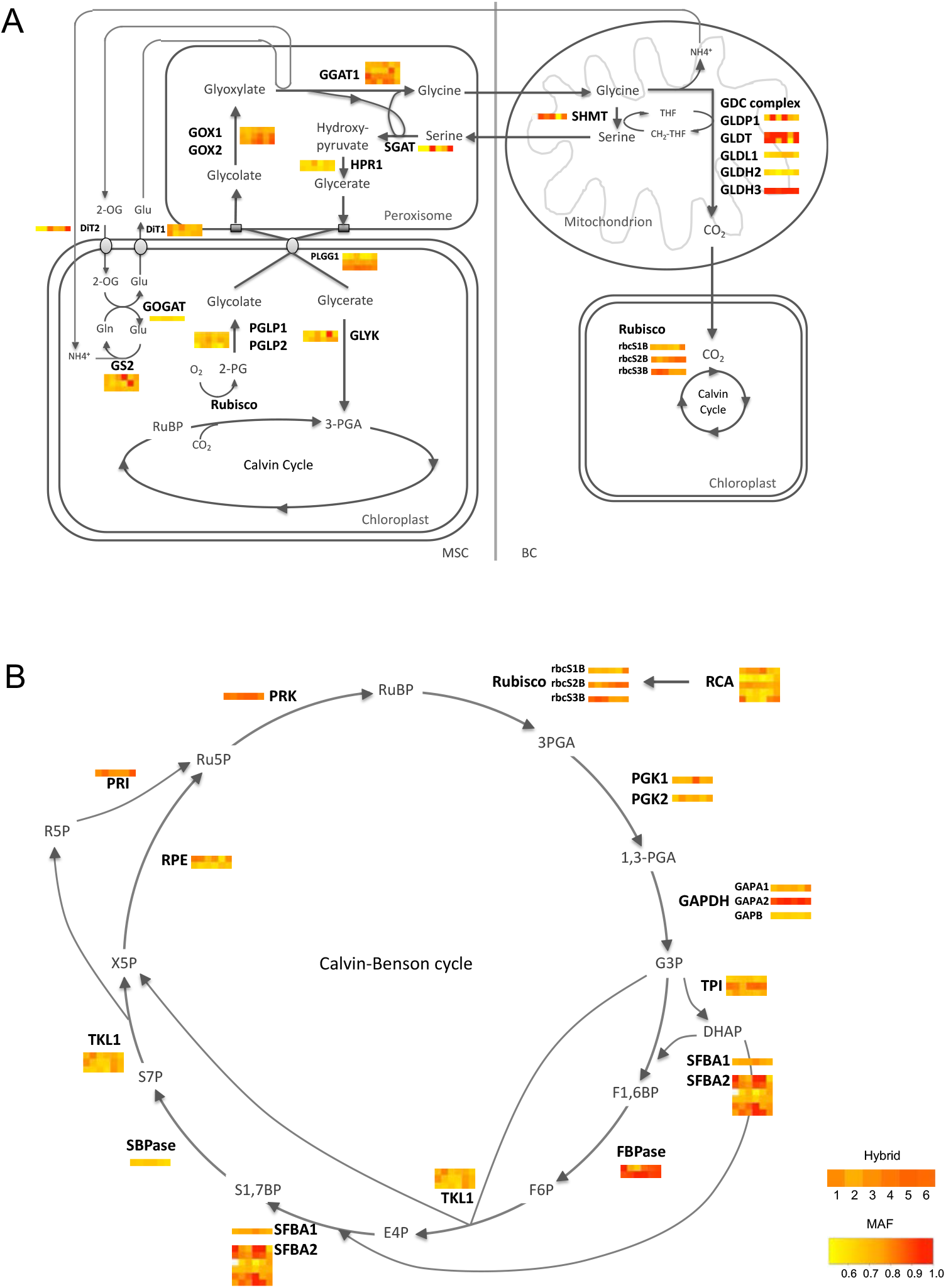
Overview on the allelic imbalance of genes involving in selected pathways of the C_3_-C_4_ glycine shuttle (A) and the Calvin-Benson cycle (B) among six *Moricandia* hybrids. The six blocks in each gene bar presented the major allele frequency (MAF) of hybrid 1 to hybrid 6 from left to right.

**Figure 7.**
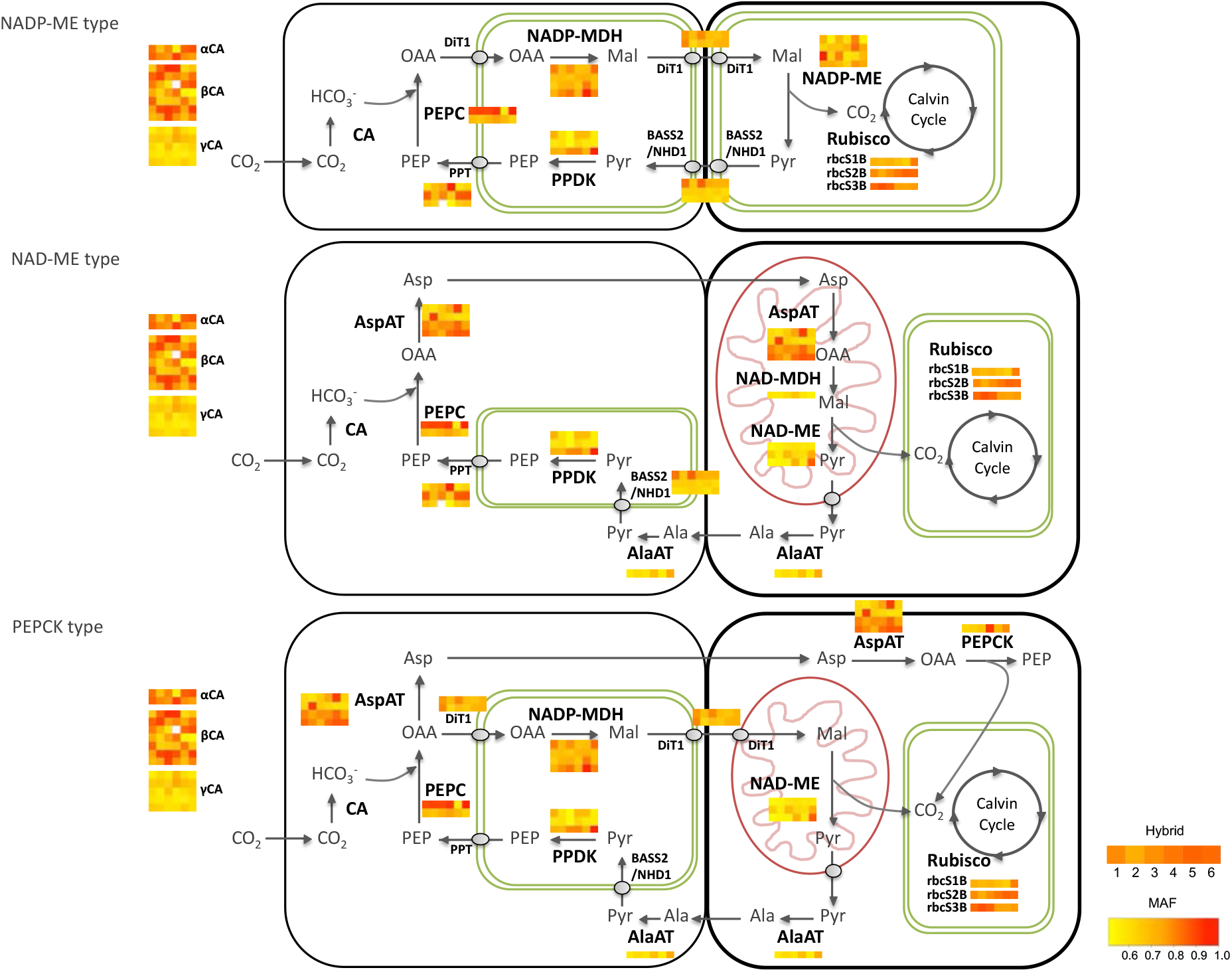
Overview on the allelic imbalance of C_4_ cycle genes among six *Moricandia* hybrids. The six blocks in each gene bar presented the major allele frequency (MAF) of hybrid 1 to hybrid 6 from left to right.

### Promoter-GUS assay on selected genes confirmed the ASE result

ASE transcripts were expected to show differences in transcriptional abundance or spatiotemporal gene expression between C_3_ and C_3_-C_4_ *Moricandia* species. To test these hypotheses, promoter-GUS assays were used. The selected genes for this assay were (1) transcripts with common *cis*-SNPs enriched in GO term chloroplast relocation (GO:0009902), and (2) ASE-transcripts with high allelic imbalance toward *M. arvensis* (MAF ≥ 0.8) across all six hybrids. GLDP1 localized exclusively to the BS cells of the leaf of C_3_-C_4_ *Moricandia* species, which is most likely regulated on transcriptional level (Adwy et al., 2019; Rawsthorne et al., 1988; Rawsthorne et al., 1988). *GLDP1* possessed *cis*-SNPs across all hybrids and an average MAF of 0.76 (Supplementary file 12; Table 2). *PHOT2* and *CHUP1* selected from the GO term chloroplast relocation had common *cis*-SNPs among six hybrids and an average MAF of 0.62 and 0.72, respectively. *DUF538* and *ATPB* were selected from transcripts with high allelic imbalance toward *M. arvensis*: *DUF538* encoding an unknown function protein showed an average MAF of 0.83; *ATPB* was annotated as ATP synthesis coupled proton transport with an average MAF of 0.96 (Supplementary file 11; Table 2).

**Table 2.**
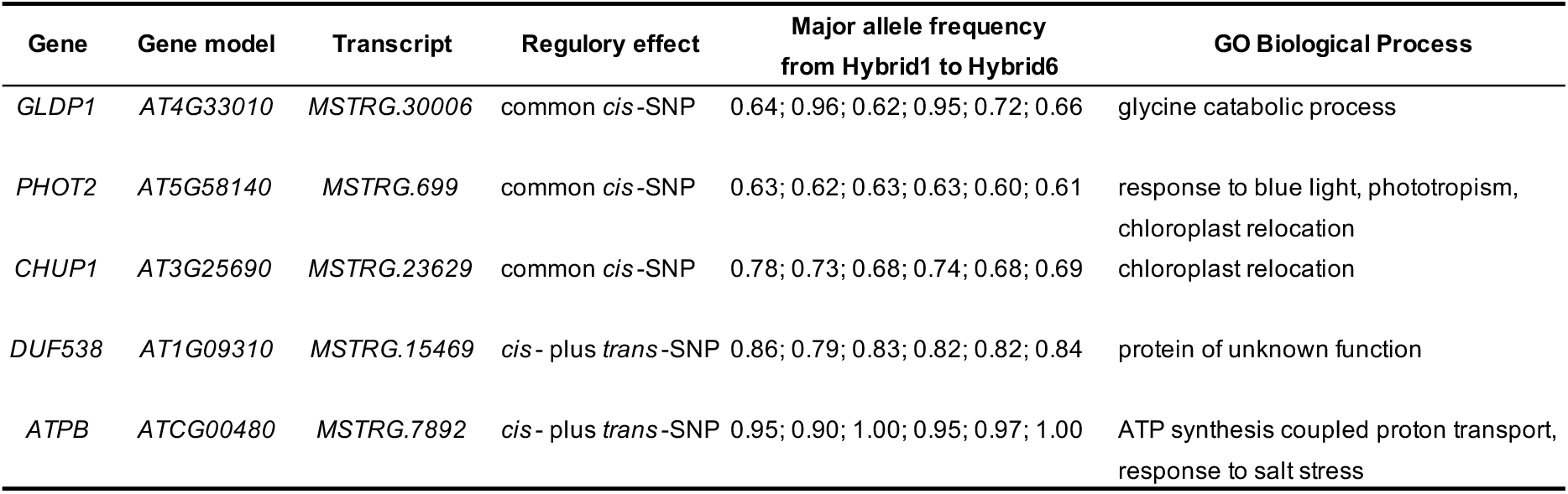
List of selected genes in *Moricandia*.

Approximately 2 kb upstream promoter region of genes from *M. arvensis* and *M. moricandioides* were amplified, fused with GUS reporter gene, and the recombinant constructs were introduced into *A. thaliana* (Supplementary file 13). GUS staining demonstrated the spatial gene expression difference between *M. arvensis* and *M. moricandioides* (Figure 8). *GLDP1*, as control, showed different cell-specific regulation between *Moricandia* species. *MaGLDP1* promoter drove GUS expression in cells surrounding veins and GUS staining of that of *M. moricandioides* (*MmGLDP1*) was observed on the whole leaf. GUS expression driven by *MaPHOT2* promoter was observed in roots and slightly in shoots of two-week-old seedlings. However, that of *MmPHOT2* was detected predominantly surrounding the leaf mid rib, trichomes in leaves, and shoots. pMaCHUP1::GUS expression was detected in the whole leaf and slightly in shoots, where GUS expression driven by *MmCHUP1* promoter was stronger in the veins and shoots. *MaDUF538* promoter drove GUS expression in parts of the leaf blade, shoots, and roots; in contrast, GUS expression driven by *MmDUF538* promoter was only detected in roots. *MaATPB* promoter resulted in vein-, root-, and shoot-preferential GUS expression veins; GUS staining of p*MmATPB*::GUS transgenic plants was detected also in leaves, but with different expression pattern between leaves. Thus, with the exception of *ATPB*, promoter-GUS assay could associate ASE on SNP- and transcript-level with specific spatial gene expression.

**Figure 8.**
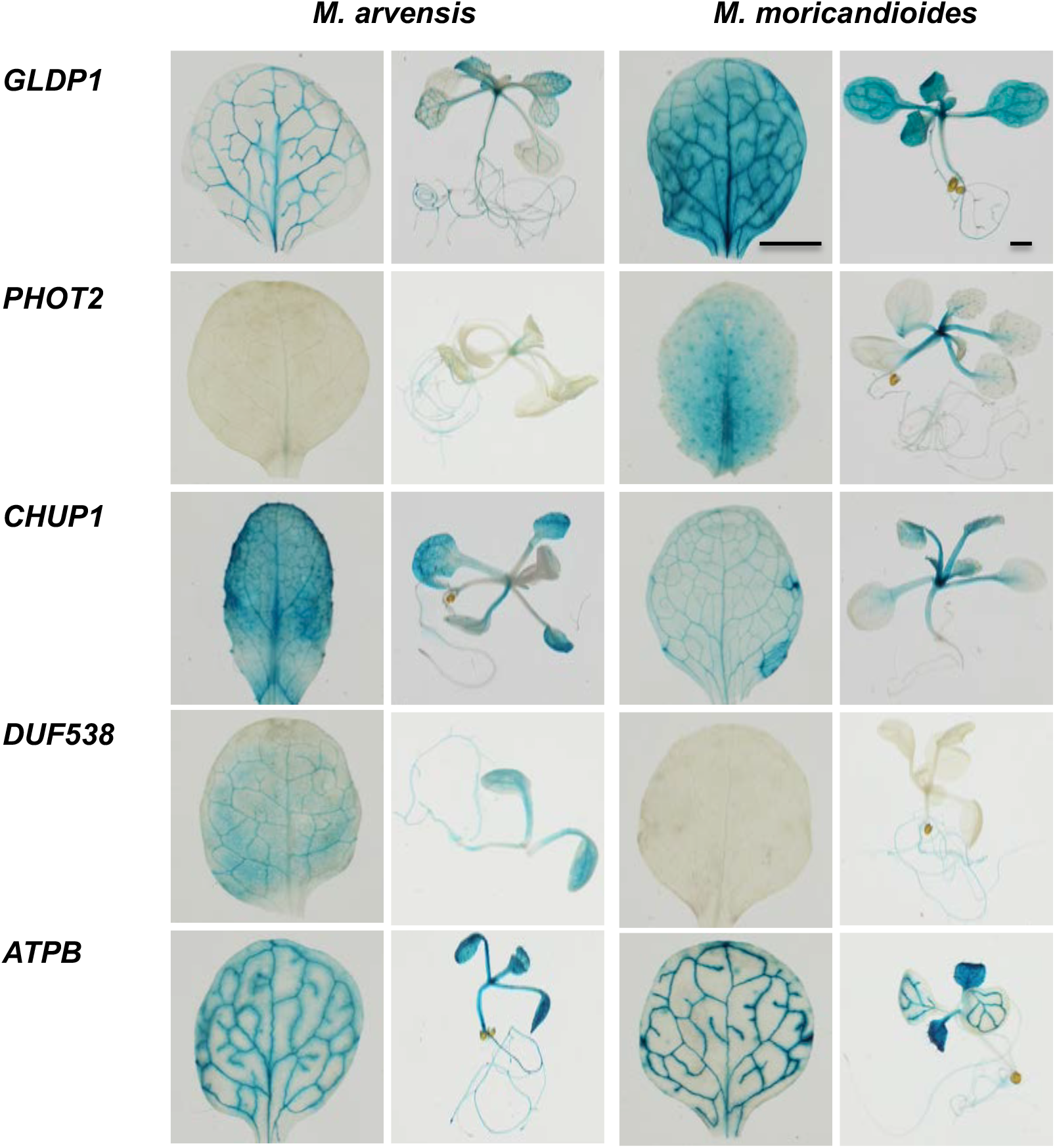
Promoter-GUS assay of *Moricandia* selected genes expressed in *A. thaliana. GLDP1* served as the positive control, possessing *cis*-SNP and defined as ASE-transcript with major allele frequency (MAF) ≥ 0.7; *PHOT2* and *CHUP1* had common *cis*-SNP among hybrids; *DUF538* and *ATPB* demonstrated strong allelic imbalance with MAF ≥ 0.8. Bar, 1 mm.

## Discussion

### ASE analysis is a powerful strategy for understanding the evolution of C_3_-C_4_ photosynthesis

The evolution of C_4_ photosynthesis via intermediate forms was achieved by regulatory diversions from the ancestral C_3_ leaf metabolism (Heckmann, 2016; Reeves et al., 2017). The regulatory mechanisms realising the C_4_ specific expression patterns have however only been studied for selected genes, mainly from the C_4_ shuttle pathway (Schlüter und Weber 2020). Our knowledge about the genetic and molecular background of C_3_-C_4_ intermediacy is so far limited to regulatory elements in the promoter of the *GLDP* gene (Adwy et al., 2019; Schulze et al., 2013).

By production of interspecific hybrids between C_3_-C_4_ *M. arvensis* and C_3_ *M. moricandioides*, novel insights into the regulatory divergences of photosynthetic systems in *Moricandia* are now obtained by ASE on SNP and transcript level, and numerous *cis*-regulatory variations between C_3_-C_4_ and C_3_ *Moricandia* species were discovered. Transcripts harboring common *cis*-SNPs dominated the major photosynthetic pathways and chloroplast relocation between C_3_-C_4_ and C_3_ *Moricandia* species.

Whole leaf transcriptome analyses of C_3_ and C_3_-C_4_ *Moricandia* species identified only very few differentially expressed genes. The *GLDP* gene, previously shown to display different spatial expression patterns in leaves from the two photosynthesis types was not flagged in whole-leave differential gene expression analysis (Schlüter et al., 2017). To the contrary, the ASE analysis reported here clearly identified *cis-*derived effects for *GLDP1* in all hybrids. Additional genes connected to the glycine shuttle (*GOX, GGAT*, other *GDC* complex subunits, *SHMT, GS2, DiT, AspAT*) contained also at least one *cis*-SNP among hybrids and demonstrated strong ASE (Supplemental Table 11; Figure 6A). Of the roughly 12,400 *Moricandia* transcripts, 27% showed evidence of ASE and 3% demonstrated extreme allelic imbalance. The results support our current hypothesis that the installation of the glycine shuttle requires regulatory adjustments in the whole photorespiratory pathway, but also adjacent pathways such as the nitrogen metabolism, Calvin Benson cycle, redox metabolism and transport processes between cellular organs as well as mesophyll and bundle sheath (Mallmann et al., 2014; Schlüter et al., 2017). Candidate genes from all these pathways showed evidence of *cis* regulation in the investigated hybrids (e.g. *CA, PEPC, PPT, NADP-MDH, DIT, NADP-ME, BASS2, PPDK, PPT, AspAT, PEPCK* and many Calvin Benson cycle genes).

Beside leaf biochemistry, the installation of an efficient glycine shuttle is based on general activation of BS metabolism, as well as BS-specific changes in organelle number and localization (Lundgren, 2020). ASE analysis identified numerous genes predicted to be involved in chloroplast movement and localization. Consistent with this observation, promoter GUS studies with two candidate genes involved in chloroplast positioning within the cell (*PHOT2* and *CHUP1*) revealed differences in their expression pattern in C_3_ and C_3_-C_4_ leaves. Chloroplast movements are important for adjustment of photosynthesis to different light conditions, the differential regulation of the two proteins in C_3_ and C_3_-C_4_ *Moricandia* species suggest they also play a role in BS specific chloroplast positioning.

The lack of knowledge about the mechanisms underpinning the C_3_-C_4_ and also C_4_ specific anatomical features represents a serious bottleneck for the engineering of more efficient photosynthetic pathways into C_3_ crop plants. ASE analysis of interspecific hybrids could help to identify key players of these anatomical changes, especially when expanded to different stages of leaf development.

### *Moricandia* interspecific hybrids showed C_3_ parental-like phenotypes

C_3_-C_4_ × C_3_ *Moricandia* hybrids were successfully created, but they were sterile most likely because of chromosome mismatching and irregular meiotic division of pollen mother cells. Similar problems were observed in trials using species in *Panicium* and *Flaveria* (Bouton et al., 1986; H. R. Brown & Bouton, 1993). However, interspecific hybrids of *A. prostrata* (C_3_) and *A. rosea* (C_4_) were fertile (Oakley et al., 2014). The six hybrids in this study demonstrated intermediate, non-uniform phenotypic characteristics between those of the parents. The CO_2_ compensation points and leaf anatomy were thereby more similar to that of the C_3_ parent (Figure 2; Table 1; Supplementary file 1 and 2). Additionally, a PCA showed that the gene expressions of *Moricandia* interspecific hybrids were also closer to that of C_3_ parents compared to that of the C_3_-C_4_ parent (Supplementary file 4B). Our finding was in accordance with intermediate phenotypic characteristics of CO_2_ compensation points and organelle quantities in BS cells observed in interspecific hybrids of C_3_-C_4_ and C_3_ *Panicum* species and intergeneric hybrids of *M. nitens* (C_3_-C_4_) × *Brassica napus* (C_3_) (Brown et al., 1985; Rawsthorne et al., 1998). The leaf anatomy of hybrids from *M. nitens* (C_3_-C_4_) × *B. napus* (C_3_) resembled that of the C_3_ parent (Rawsthorne et al., 1988). The increase in C_3_-C_4_ proportion also increased the intermediate phenotype. The dominance of C_3_-C_4_ phenotypes (CO_2_ exchange and confined GDC in BS cells) increased with the C_3_-C_4_ genome constitution in *D. tenuifolia* (C_3_-C_4_) × *R. sativus* (C_3_) hybrids and *M. arvensis* (C_3_-C_4_) × *B. oleracea* (C_3_) and their reciprocal crosses (Ueno et al., 2003, 2007), and the same phenomenon was found in backcrosses of *B. alboglabra* (C_3_) × *M. arvensis* (C_3_-C_4_) to the C_3_-C_4_ parent (Apel et al., 1984). These findings, together with our observation, suggest that genes regulating C_3_-C_4_ characteristics showed additive effects in C_3_-C_4_ × C_3_ hybrids. For the *GLDP* gene, the C_3_ copy for instance could be responsible for almost C_3_-like expression in the M. Thereby, *Moricandia* C_3_-C_4_ × C_3_ hybrids more resembled C_3_ parent, because C_3_-C_4_ species had also the C_3_ genetic background.

### *Cis* mechanisms play a major role in evolution of C_3_-C_4_ intermediacy

*Cis*-regulatory effects presented larger impacts than *trans*-acting divergences on *Moricandia* interspecific hybrids: 36% and 6% of assayed SNPs were discovered as *cis*-SNP and *trans*-SNP, respectively. Our results are in accordance with previous findings showing that *cis*-regulatory changes are more prevalent in interspecific hybrids (long evolutionary time-scales), whereas *trans*-regulatory divergence is more frequently observed in intraspecific hybrids (short evolutionary time-scales) (McManus et al., 2010; Rhoné et al., 2017; Stern & Orgogozo, 2008). The dominance of *cis*-regulatory divergence was also shown in ASE studies on interspecific hybrids, such as in *Atriplex* and poplar (Sultmanis, 2018; Zhuang & Adams, 2007). The relative proportions of regulatory effects were similar in all six Ma×Mm hybrids, but many of them were sample-specific. Only 8.7% of *cis*-SNPs and 1% of *trans*-SNPs were found consistently across all interspecific hybrids. It was also described in the literature that ASE genes are often unique in different hybrids (Lemmon et al., 2014; Steige et al., 2015). This finding might explain that the hybrids didn’t show uniform phenotypic characteristics, and together with the sterility of hybrids, it supports the problems with different chromosome arrangement between the parents.

*cis*- and *trans*-regulatory divergences have different impacts on the inheritance and evolution of gene expression, and hence also on different biological processes (McManus et al., 2010; Meiklejohn et al., 2014). This is related to the genetic nature of transcriptional regulations: *cis*-regulatory sequences, located in promoter regions, UTRs, and introns, modulate the binding of *trans*-acting factors to DNA, therefore affecting the transcription of nearby genes, whereas *trans*-element, such as transcription factors and long noncoding RNA, are able to affect the expression of many genes (Wray, 2007). In our study, ASE would detect regulatory differences in all affected pathways. GO and pathway enrichment analysis on transcripts with common *trans*-SNPs indicated very general biological pathways. Transcripts with common *cis*-SNPs on the other hand showed enrichment in major photosynthetic pathways and chloroplast relocation, thus indicating that the differences in the parental photosynthesis types were *cis*-regulated. For instance, transcripts with common *cis*-SNP were overrepresented in isopentenyl diphosphate biosynthesis. In higher plants, the formation of isopentenyl diphosphate, the central intermediate of all isoprenoids, was compartmentalized, such as sterols in cytosol or carotenoid, phytol, and chlorophyll in plastids (Lange and Croteau, 1999), which is likely regulated by *cis*-acting divergences. In the present study, *cis*-specificity accumulated in the gene category of chloroplast relocation and anatomical structure morphogenesis in *Moricandia*. The anatomical adjustment, such as centripetal accumulation of mitochondria and chloroplasts in BS cells, was a prominent characteristic for reduced compensation points in C_3_-C_4_, resulting from the glycine shuttle (Schlüter et al., 2017). Regulatory divergences were for instance found for two genes responsible for chloroplast localization within the cells. *PHOT2* mediates blue light responses and is involved in phototropism, chloroplast movement, stomatal opening, leaf development and photosynthetic efficiency (Hart et al., 2019). *CHUP1* is involved in chloroplast positioning within the cell and interaction with the actin cytoskeleton (Oikawa et al., 2008). The expression of both genes was tested by promoter-GUS assay in Arabidopsis and different patterns could be observed for the Ma and Mm derived promoters. Of course, elements in untranslated gene regions (5’UTR and 3’UTR), coding regions, and introns have been reported to affect steady-state transcript amounts could be active in addition to the promoter region (Barrett et al., 2012; Hernandez-Garcia and Finer, 2014).

*cis*-regulatory divergences are additive in heterozygotes, which were visible and preferentially accumulated during evolution, therefore playing a crucial role in evolution of adaptive traits (Lemos et al., 2008; Meiklejohn et al., 2014; Wittkopp et al., 2008). In our study, transcripts with *cis*-specificity showed preferential expression toward C_3_-C_4_ *M. arvensis* alleles in hybrids (Supplementary file 10), although the phenotype of hybrids more resembled that of the C_3_ parent. During evolution of the selfing syndrome in *Capsella*, alleles from the self-compatible species *C. rubella* were expressed at higher level than the outcrossing *C. rubella* alleles in their hybrids (Steige et al., 2015). *cis*-regulatory divergences dominated the positive selection and the adaptive improvement during maize domestication from teosinte, and genes with *cis*-regulatory effect demonstrated a directional expression bias toward maize (Lemmon et al., 2014). An expression preference was however not found in C_3_ × C_4_ *Atriplex* hybrids (Sultmanis, 2018). Transcripts with *cis*-specificity showing a directional expression bias toward *M. arvensis* were more abundant in GO terms, involving in anatomical structure morphogenesis, transmembrane transporter activity and localization, compared to transcripts expressed biased to *M. moricandioides*. The installation of the glycine shuttle could therefore be associated with *cis*-mediated upregulation of genes involved in leaf anatomy and transport activities.

Our study suggests that many *cis*-regulatory effects, favored in adaptive phenotypic traits during evolution, were additive in C_3_-C_4_ × C_3_ *Moricandia* hybrids. This is consistent with the previously predicted long and smooth path of C_4_ photosynthesis evolution (Heckmann et al., 2013; Williams et al., 2013).

## Conclusion

Interspecific hybrids between C_3_ and C_3_-C_4_ *Moricandia* species possessed phenotypic characteristics of CO_2_ compensation point and chloroplast accumulation in BS cells between those of the parental species, however more resembling those of the C_3_ parent. We showed that *cis*-regulatory divergences have a large impact on *Moricandia* interspecific hybrids, and the corresponding transcripts were found to be enriched in major photosynthetic pathways and chloroplast relocation. We further observed that *cis*-specificity caused enhanced expression of C_3_-C_4_ alleles in categories such as anatomical structure morphogenesis and transmembrane transporter activity. *cis* mechanisms contributed to the installation of the glycine shuttle, playing an important role in the early evolutionary steps of C_4_ photosynthesis. With the genetic information of parental species, the RNA-Seq dataset and ASE approaches, we investigated *cis*- and *trans*-acting divergences on a transcriptome-wide scale, which helps us to understand the role of transcriptional regulations during evolution of C_4_ photosynthesis. It also provides possible targets for engineering C_3_-C_4_ characteristics into C_3_ plants.

## Materials and Methods

### Plant materials

Seeds from *Moricandia* were surface-sterilized using chloride gas and germinated on half MS medium for one week. The seedlings were then transferred individually to pots with soil and grown in the growth chamber under 12h/12h light/dark conditions with 23°C/20°C day/night. The anthers of *M. arvensis* (IPK Gatersleben: MOR1, a C_3_-C_4_ intermediate, as maternal plant) were removed and their stigma was bagged one day before the artificial cross-pollination. The pollen from *M. moricandioides* (Botanical Garden Osnabrück: 04-0393-10-00, a C_3_ species, as paternal plant) were collected and transferred to the receptive stigma of *M. arvensis*. The reciprocal crosses were done in the same way. The seeds from parents and hybrids germinated and grew following the same procedure described before. Two-week-old leaves of parents and hybrids were used for DNA extraction for genotyping and for promoter region amplification. The two youngest leaves from four-week-old plants were collected as materials for RNA-Seq. Moreover, the mature rosette leaves were taken for leaf anatomy and gas exchange analysis. Seeds from *Arabidopsis thaliana* wild-type plants ecotype Col-0 and the transgenic lines were surface sterilized by vapor phase seed sterilization, further germinated on half MS medium with cold treatment for two days in the dark, and then transferred to the growth chamber under 10h/14h light/dark conditions with 22°C/20°C day/night for 10 days. The seedlings were later transferred individually to pots with soil and grown in the growth chamber. The two-week-old Arabidopsis transgenic plants and the wild-type plants were collected for further GUS staining analysis.

### Leaf anatomy

The 2 mm^2^ leaf sections were taken near the midrib of the top third of mature rosette leaves for the leaf ultrastructural analysis. For araldite embedding, leaf sections were fixed with fixation buffer (2% paraformaldehyde, 2% glutaraldehyde), dehydrated by an acetone dilution series, and embedded with an araldite series according to the protocol (Fineran & Bullock, 1972) with modifications. The sections were transferred to the mold filled with fresh araldite and polymerized at 65°C for two days. Semi-thin sections in 2.5 μm thickness obtained by cutting with a glass knife were mounted on slides, stained with 1% toluidine blue for 2 min and washed by distilled water. The leaf ultrastructure was examined under the light microscope, Zeiss Axiophot microscope (Carl Zeiss Microscopy GmbH, Göttingen, Germany).

### Photosynthetic gas exchange

The mature rosette leaves were chosen to measure gas exchange characteristics using a LI-6400XT Portable Photosynthesis System (LI-COR Biosciences, Lincoln, USA) with the settings according to the manufacturer’s instructions with modifications: the flow of 300 μmol s−1, the light source of 1500 μmol m−2 s−1, the leaf temperature of 25°C, and the vapor pressure deficit based on leaf temp less than 1.5 kPa. The CO_2_ response curve, the so called A-Ci curve, was captured by detecting net CO_2_ assimilation rates under different intercellular CO_2_ concentrations. A partial A-Ci curve obtained with measurements at 400, 100, 80, 65, 45, 25, 15, and 400 ppm CO_2_ was used to calculate the CO_2_ compensation points of parental genotypes and hybrids.

### Sample preparation and RNA sequencing

We selected 12 plants including three replicates of *M. arvensis*, three plants of *M. moricandioides* and six F_1_ interspecific hybrids (Ma×Mm). The six hybrids demonstrated relative lower CO_2_ compensation points among hybrids and differed in leaf anatomy patterns (Figure 2; Table 1). Compared to the parents, hybrids 1, 2, 5, 6 had fewer organelles in BS cells, whereas hybrid 3 and 4 showed very little organelle in BS cells (Supplementary file 1). Total RNA of parental species and interspecific F_1_s was extracted using the RNeasy Plant Mini Kit (Qiagen, Hilden, Germany). Then, 17 μl total RNA (100 ng/μl) was added with 2 μl buffer and 0.5 μl RNase-free DNaseI enzyme (New England Biolabs GmbH, Frankfurt am Main, Germany) incubating on ice for 30 s. The treatment was stopped by adding 2 μl 50 mM EDTA and incubated at 65°C for 10 min. The quality of RNA and DNaseI treated RNA was assessed on a Bioanalyzer 2100 (Agilent, Santa Clara, USA) with an RNA Integrity Number (RIN) value ≥ 8. Subsequently, cDNA libraries were prepared using 1 μg of total RNA with the TruSeq RNA Sample Preparation Kit (Illumina, San Diego, USA). The cDNA library was qualified on the Agilent Technologies 2100 Bioanalyzer to check the library quality and fragment size of the sample. RNA-Seq was performed on an Illumina HiSeq 3000 platform at the BMFZ (Biologisch-Medizinisches Forschungszentrum) of the Heinrich-Heine University (Düsseldorf, Germany) to gain 150 bp paired-end reads. In total, we obtained 54.97 Gb of RNA-Seq data, with an average of 4.58 Gb per sample. On average, 27 million reads per library were obtained. The sequencing quality was examined using FastQC v.0.11.5. Quality scores across all bases were generally good but showed lower quality at the end of reads observed in few samples.

### Read mapping and variant calling

In our study, the C_3_ *M. moricandioides* genome assembly was chosen as the draft reference genome, because it possesses a higher quality over C_3_-C_4_ *M. arvensis* by having more reliable number of repetitive elements. Mapping all the RNA-Seq reads to *M. moricandioides* might lead to underestimation of the transcript level of *M. arvensis* allele, however it has no impact on comparing the allele ratio between parents and hybrids as well as transcripts dominated by *M. arvensis* allele. The RNA-Seq reads were mapped on a draft reference genome of *M. moricandioides* (Lin et al., 2021) using STAR v.2.5.2b (Dobin et al., 2013). After duplication marking, base quality recalibration, we used a simulated set of SNPs as known variants for preparing analysis-ready RNA-Seq reads. The variant calling was conducted according to the Genome Analysis Toolkit (GATK) best practice (DePristo et al., 2011). Variant discovery was performed jointly the three *M. arvensis* replicates using the UnifiedGenotyper with *M. moricandioides* as the reference, which output a raw vcf file containing 1,748,436 SNP callings. The variant calling vcf file and aligned RNA-Seq reads were further input into ASEReadCounter from GATK to obtain read counts at each SNP site. Only SNP sites with more than 20 total read counts of parental species and less than 3 counts at the other parental species were processed for further allele specific expression (ASE) analysis.

### Differential gene expression analysis

Transcriptome comparison between species was performed with the DESeq2 tool (Love et al., 2014) in R (www.R-project.org) using the Benjamini–Hochberg (BH) adjusted false discovery rate ≤ 0.01 as the cut-off for significant differential expression (Benjamini & Hochberg, 1995). The Chi-square test was applied to test the overrepresentation of upregulated transcripts in selected pathways, such as glycine shuttle, C_4_ cycles, Calvin-Benson cycle, and mitochondrial e^-^ transport. The principle component analysis (PCA) analysis was performed on the rlog-transformed gene expression data to explore and visualize distances between samples.

### Allele specific expression (ASE) analysis

In hybrids, parental alleles were expressed under the same genetic background which made it possible to distinguish between *cis*- or *trans*-regulatory effects by calculating and comparing the allele ratio of the parents (A: PA1/PA2) and that of hybrids (B: F1A1/F1A2). A binomial test was applied to test if F1A1 is equal to F1A2 using a *P*-value adjusted by the BH procedure. On the other hand, Fischer’s exact test was used to assess the significant difference between the ratio of parental alleles (PA1/PA2) and the allele ratio of hybrids (F1A1/F1A2) where the *P*-value was adjusted by the BH method. Four regulatory effects were defined according to the following conditions: *cis-* only, B≠1 and A=B; *trans-* only, B=1 and A≠B; *cis*-plus *trans*-, B≠1 and A≠B; no *cis*-no *trans*-, B=1 and A=B. ASE analysis on SNP-level was conducted on six hybrids individually on a set of 120,200 SNPs, demonstrating polymorphisms on 14,004 transcripts.

To integrate SNP information into a gene-specific measure, R package, meta-analysis-based allele-specific expression detection (MBASED) was applied to measure the allelic imbalance (Mayba et al., 2014). MBASED applied the principles of meta-analysis on combining the information of every SNP site within a single transcript (a single unit of expression) in the absence of the prior information of phased data, the genetic information of hybrids. The ASE was evaluated based on the transcripts with at least one SNP site. The pseudo-phasing based “major” haplotype of genes took the allele with higher counts as the major allele, resulting in the higher estimates of the major allele frequency (MAF, ranging from 0.5 to 1.0). At least 10^6^ simulations were carried out to obtain appropriate assessment of the statistical significance.

### ASE verification by qPCR

A quantitative real-time PCR (qPCR) assay using SNP-specific primers for four selected genes was used to validate the RNA-Seq and ASE results (Supplementary file 14). The DNaseI treated RNA was preceded to cDNA synthesis according to the manufacturer’s instructions with modifications. First, a total of 1 μg RNA was mixed with 1 μl oligo-dT primer, 10 mM dNTP-Mix, 4 μl 5X Firstrand-Buffer, and 2 μl 0.1 M DTT and incubated at 42°C for 2 min. The mix was combined with 1 μl Invitrogen SuperScript™ II Reverse Transcriptase and incubated at 42°C for 50 min for cDNA synthesis. The heat inactivation of reverse transcripts was conducted with incubation for 15 min at 70°C. SNPs on *Moricandia* orthologs of *GLDP1, ASP3, γCA2, PPA2* were chosen for designing SNP-specific qPCR primers (Supplementary file 15). The *Moricandia* ortholog (*MSTRG*.*23175*) of Arabidopsis housekeeping gene Helicase (*AT1G58050*) was tested and selected as reference housekeeping gene. The qPCR amplification was carried out in a total reaction of 20 μL containing 0.5 μL forward primer (10 ng/μL), 0.5 μL forward primer (10 ng/μL), 5 μL 5 ng/μL cDNA template, 4 μL ddH_2_O, and 10 μL SYBR® Green qPCR SuperMix (Thermo Fisher Scientific, Schwerte, Germany). The qPCR reaction was carried out on StepOne™ Real-Time PCR System (Applied Biosystems™, Waltham, USA) by following program: initial denaturation at 95 °C for 60 s, 40 cycles of denaturation at 95 °C for 15 s and annealing at 60 °C for 30 s. The delta CT value was calculated by normalized sample’s CT value with that of the housekeeping gene.

### Transcriptome annotation

After comparing the *M. moricandioides* predicted protein from TransDecoder (Haas et al., 2013) to UniProtKB (both Swiss-Prot and TrEMBL,ed on April 3, 2019) (Camacho et al., 2009) using BLASTP (UniProt, 2019) with e-value < 1e-5, we summarized the functional annotation in the form of “Human Readable Description” by the AHRD pipeline (https://github.com/groupschoof/AHRD). Then, to determine the phylogenetic relationships among *M. arvensis* and *M. moricandioides*, the predicted protein sequences of them together with *A. thaliana* were applied to OrthoFinder v.2.3.3 (Emms & Kelly, 2019).

### Biased transcript with *cis*-specificity

The transcripts with *cis*-specificity (*cis*-SNPs or *cis plus trans*-SNPs) were classified to two categories based on the gene expression direction in hybrids: transcripts with biased expression toward *M. arvensis* (Ma-biased) and toward *M. moricandioides* (Mm-biased). The biased transcripts were annotated with corresponding GO terms derived from *A. thaliana*. Afterward, the gene ontology comparison between Ma-biased and Mm-biased transcripts were conducted on WEGO 2.0 website (Ye J et al., 2018) and visualized using R.

### Gene ontology term and pathway enrichment analysis

To recognize the function of transcripts with *cis*-SNPs (common *cis*-SNPs) and with *trans*-SNPs (common *trans*-SNPs) shared by all studied hybrids, a custom mapping file was created, containing *Moricandia* transcript names and the corresponding GO terms derived from *A. thaliana* genes. The 2,236 and 45 *Moricandia* transcripts with common *cis-*SNPs and common *trans*-SNPs, respectively, were processed with the custom mapping file by topGO R-package for gene set enrichment analysis of biological processes (Alexa et al., 2006).

Pathway enrichment analysis was conducted on the mRNA sequences of 2,236 and 45 *Moricandia* transcripts with common *cis-*SNPs and *trans*-SNPs, respectively, with the KEGG Orthology Based Annotation System (KOBAS) (Xie et al., 2011). A BH adjusted false discovery rate of 0.05 served as the threshold to define the significantly enriched pathways (Benjamini & Hochberg, 1995).

### Promoter-GUS assay and plant transformation

The 5’ upstream regions of the *GLDP1, CHUP1, DUF538, ATPB* genes of *M. arvensis* and *M. moricandioides* were fused to the GUS reporter gene in the binary plant vector pCambia1381. The primers for amplifying the promoter region were included a BamHI site at the 5’ border and a NcoI site at the 3’ end of the DNA fragment. The DNA fragment was inserted into pCambia1381 by homologous recombination using the Gibson Assembly Cloning kit (New England Biolabs, catalog number: E5510S). The predicted promoter region of the *PHOT2* gene of *M. arvensis* and *M. moricandioides* was cloned to a Gateway donor vector pDONR207, and then further cloned to a Gateway destination vector pGWB3, which was for C-terminal GUS fusions. The primers for amplifying the promoter region of *PHOT2* gene were included an attB1 sequence at the 5’ border and an attB2 sequence at the 3’ end of the DNA fragment. The +1 positions of the candidate genes were defined in different ways, shown in Supplemental Table 13. All generated constructs were verified by colony-PCR and DNA sequencing.

The promoter-GUS constructs were transformed into *Agrobacterium tumefaciens* strain GV310::pMP90 (Koncz and Schell, 1986) by electroporation. All constructs were verified again by colony-PCR and DNA sequencing. The *Agrobacterium* introduced with the promoter-GUS constructs were transformed in four to six-week-old *A. thaliana* (col-0) by floral-dip method (Clough and Bent, 1998). The transformed T_1_ seeds were collected in four to six weeks after transformation, and then selected on Hygromycin B contained half MS plates for two weeks. The survival T_1_ lines were further transferred to pots with soil and verified the insertion of T-DNA by PCR.

Primers used in promoter region amplification and colony-PCR were shown in Supplementary file 16.

### GUS staining

Two to four-week-old T_1_ leaves were stained with GUS staining solution (100 mM Na_2_HPO_4_, 100 mM NaH_2_PO_4_, 1 mM Potassium-Ferricyanide K_4_[Fe(N_6_)], 1 mM Potassium-Ferrocyanide K_3_[Fe(N_6_)], 0.2% Triton X-100, 2mM X-Gluc) and incubated at 37°C in the dark for 2 to 72 hours. The GUS stained leaves were further fixed by the fixation solution (50% Ethanol, 5% Glacial acetic acid, 3.7% Formaldehyde) at 65°C for 10 min. Then, leaves were incubated in 80% Ethanol at room temperature in order to remove the chlorophyll.

## Accession numbers

Sequencing read data have been deposited at the European Nucleotide Archive under the project number PRJEB39765 (https://www.ebi.ac.uk/ena/submit/sra/#studies).

## Acknowledgements

We thank Dr. Otho Mantegazza and Nils Koppers for their bioinformatics support, and Samantha Flachbart for cDNA library preparation. Many thanks to Prof. Dr. Miltos Tsiantis for constructive suggestions and to Prof. Juliette de Meaux for encouraging us to pursue an ASE approach. We thank the gardeners for taking care of the plants and Steffen Köhler for photography support.

## Additional information

### Funding

This work was funded by grants of the Deutsche Forschungsgemeinschaft to APMW under Germany’s Excellence Strategy EXC-2048/1, Project ID 390686111, and by ERA-CAPS project C4BREED (WE 2231/20-1), and by a graduate fellowship of the International Max Planck Research School on “Understanding Complex Plant Traits using Computational and Evolutionary Approaches” to MY.L.

### Author contributions

**MY.L**. designed and performed all the experimental works and data analysis and wrote the manuscript.

**B. S**. participated in drafting the manuscript.

**U.S**. and **A.P.M.W**. conceptualized the study, supervised the experimental design and participated in drafting the manuscript.

## Additional files

### Major datasets

The following datasets were generated:

| Author(s) | Year | Dataset title | Dataset URL | Database and identifier |
| --- | --- | --- | --- | --- |
| Lin M-Y, Schlüter U, Stich B, Weber AP | 2021 | M. arvensis_plant2_RNA-Seq reads | https://www.ebi.ac.uk/ena/browser/view/ERS5424638 | Publicly available at ENA, ERS5424638 |
| Lin M-Y, Schlüter U, Stich B, Weber AP | 2021 | M. arvensis_plant5_RNA-Seq reads | https://www.ebi.ac.uk/ena/browser/view/ERS5424639 | Publicly available at ENA, ERS5424639 |
| Lin M-Y, Schlüter U, Stich B, Weber AP | 2021 | M. arvensis_plant6_RNA-Seq reads | https://www.ebi.ac.uk/ena/browser/view/ERS5424640 | Publicly available at ENA, ERS5424640 |
| Lin M-Y, Schlüter U, Stich B, Weber AP | 2021 | M. moricandioides_plant1_RNA-Seq reads | https://www.ebi.ac.uk/ena/browser/view/ERS5424641 | Publicly available at ENA, ERS5424641 |
| Lin M-Y, Schlüter U, Stich B, Weber AP | 2021 | M. moricandioides_plant7_RNA-Seq reads | https://www.ebi.ac.uk/ena/browser/view/ERS5424642 | Publicly available at ENA, ERS5424642 |
| Lin M-Y, Schlüter U, Stich B, Weber AP | 2021 | M. moricandioides_plant8_RNA-Seq reads | https://www.ebi.ac.uk/ena/browser/view/ERS5424643 | Publicly available at ENA, ERS5424643 |
| Lin M-Y, Schlüter U, Stich B, Weber AP | 2021 | M. arvensis x M. moricandioides_Hybrid1_RNA-Seq reads | https://www.ebi.ac.uk/ena/browser/view/ERS5455027 | Publicly available at ENA, ERS5455027 |
| Lin M-Y, Schlüter U, Stich B, Weber AP | 2021 | M. arvensis x M. moricandioides_Hybrid2_RNA-Seq reads | https://www.ebi.ac.uk/ena/browser/view/ERS5455028 | Publicly available at ENA, ERS5455028 |
| Lin M-Y, Schlüter U, Stich B, Weber AP | 2021 | M. arvensis x M. moricandioides_Hybrid3_RNA-Seq reads | https://www.ebi.ac.uk/ena/browser/view/ERS5455029 | Publicly available at ENA, ERS5455029 |
| Lin M-Y, Schlüter U, Stich B, Weber AP | 2021 | M. arvensis x M. moricandioides_Hybrid4_RNA-Seq reads | https://www.ebi.ac.uk/ena/browser/view/ERS5455030 | Publicly available at ENA, ERS5455030 |
| Lin M-Y, Schlüter U, Stich B, Weber AP | 2021 | M. arvensis x M. moricandioides_Hybrid5_RNA-Seq reads | https://www.ebi.ac.uk/ena/browser/view/ERS5455031 | Publicly available at ENA, ERS5455031 |
| Lin M-Y, Schlüter U, Stich B, Weber AP | 2021 | M. arvensis x M. moricandioides_Hybrid6_RNA-Seq reads | https://www.ebi.ac.uk/ena/browser/view/ERS5455032 | Publicly available at ENA, ERS5455032 |
| Lin M-Y, Schlüter U, Stich B, Weber AP | 2021 | Transcriptome assembly of M. moricandioides | https://www.ebi.ac.uk/ena/browser/view/ERS5472842 | Publicly available at ENA, ERS5472842 |

## Supplementary files

**Supplementary file 1**. Leaf micrographs of transverse sections of *M. arvensis, M. moricandioides* and their interspecific hybrids a, *M. arvensis*; b, *M. moricandioides*; c-h, hybrids 1-6. Arrow, chloroplasts. Bar, 100 µm.

**Supplementary file 2**. Leaf venations of *M. arvensis, M. moricandioides* and their interspecific hybrids a, *M. arvensis*; b, *M. moricandioides*; c-h, hybrid 1-6. Bar, 500 µm.

**Supplementary file 3**. The pollen activity test of *M. arvensis, M. moricandioides* and their interspecific hybrids dyed by Alexander staining method (Alexander, 1969) Pollens from hybrid lines were stained red, but demonstrated abnormal shapes compared to parents’ round pollens. Aborted pollen grains are stained blue-green, and non-aborted pollen grains are stained magenta-red. Bar, 20 µm.

**Supplementary file 4**. Principal component analysis of rlog-transformed gene expression data A) parental species, *M. arvensis* (Ma) and *M. moricandioides* (Mm) and B) parental species and their interspecific hybrids.

**Supplementary file 5**. GO analysis on Ma-upregulated transcripts and Ma-downregulated transcripts using topGO.

**Supplementary file 6**. Transcriptional changes in selected pathways. The heatmap indicated the log2-fold changes in transcript level of C_3_–C_4_ species *M. arvensis* compared to the C_3_ species *M. moricandioides*. Blue and red indicates reduced and enhanced transcript abundance in C_3_–C_4_, respectively. *, adjusted P-value < 0.05; **, adjusted P-value < 0.01. Bold, transcripts with the highest expression among isoforms.

**Supplementary file 7**. List of common SNPs and transcripts harboring common SNP.

**Supplementary file 8**. GO analysis on common *cis*-SNPs and common *trans*-SNPs using topGO.

**Supplementary file 9**. Significantly enriched pathways identified in transcripts with common *cis*-SNPs and common *trans*-SNPs using KOBAS database.

**Supplementary file 10**. Number of biased transcripts with *cis*-specificity among six hybrids.

**Supplementary file 11**. List of transcripts showed extreme allelic imbalance with major allele frequency ≥ 0.9 in all hybrids.

**Supplementary file 12**. Enrichment of regulatory effects in selected pathways. 0, no *cis*-SNP; 1, at least one *cis*-SNP found in hybrid line; 2, common *cis*-SNP among hybrids.

**Supplementary file 13**. Selected gene list for promoter-GUS assay.

**Supplementary file 14**. Confirmation of RNA-Seq data by allele-specific RT-PCR of *M. arvensis* × *M. moricandioides* hybrid.

**Supplementary file 15**. qPCR primer list for ASE verification.

**Supplementary file 16**. Primer list for promoter-GUS assay.

## Supplementary method

### Pollen activity assay

The pollen viability of *M. arvensis, M. moricandioides*, and their hybrids was observed followed modified Alexander’s staining method (Alexander, 1969). The primary inflorescences with mature pollens were collected one day after flowering and then incubated in 1:50 staining solution for 5 min. The stock solution was comprised of 10 ml 96% ethanol, 1 ml 1% malachite green (w/v, in 96% ethanol), 25 ml glycerol, 5 ml 1% acid fuchsin (w/v, in dH_2_O), 4 ml glacial acetic acid, and 100 ml dH_2_O. The phase contrast images of dyed pollens were obtained under inverted microscopy (Eclipse Ti, Nikon).

## Notes

### Competing Interest Statement

The authors have declared no competing interest.

