## Supplementary Figures and Tables for "*Cis*-regulatory divergence underpins the evolution of C_3_-C_4_ intermediate photosynthesis in *Moricandia*"

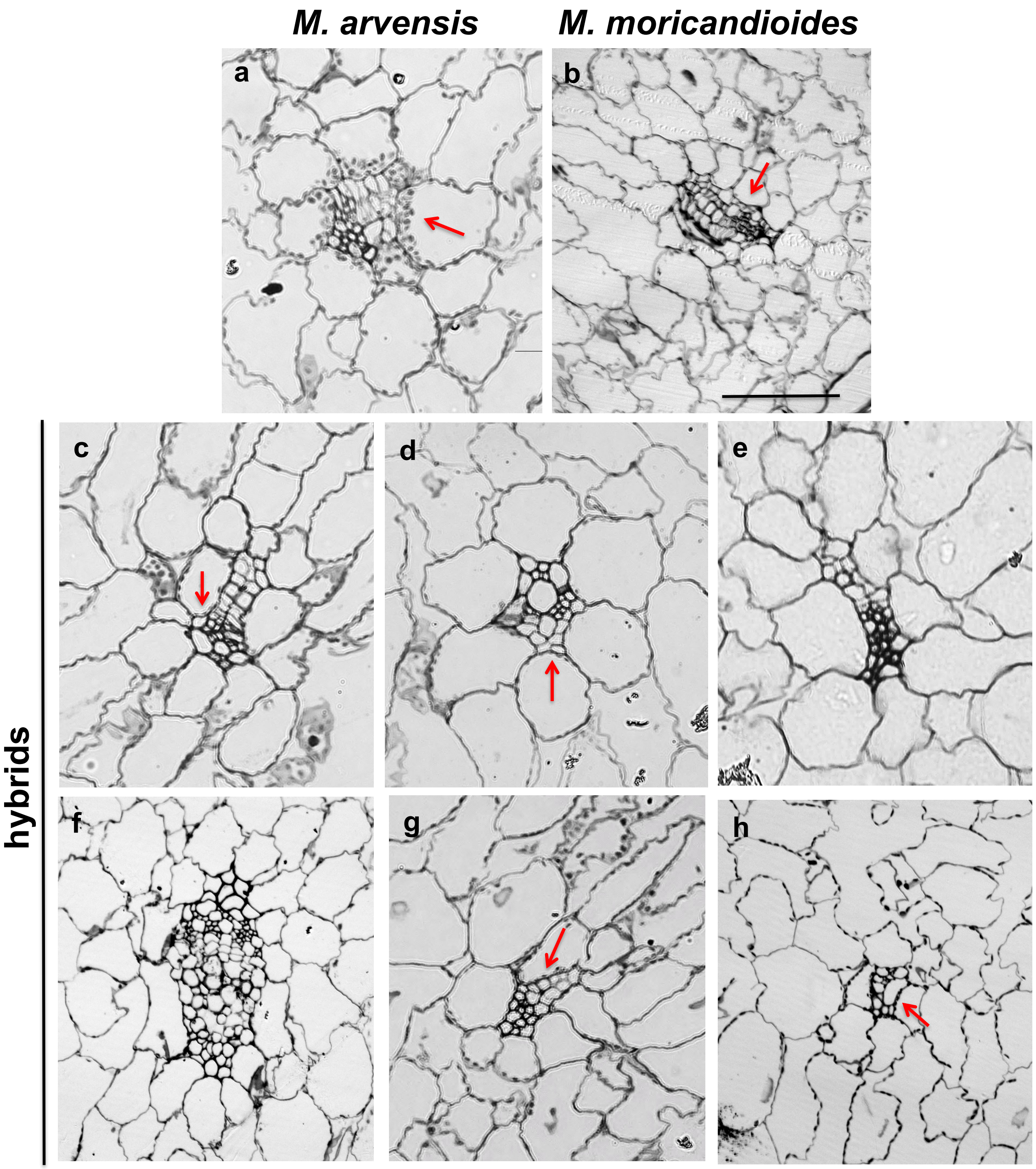


**Supplementary file 1.** Leaf micrographs of transverse sections of *M. arvensis*, *M. moricandioides* and their interspecific hybrids. a, *M. arvensis*; b, *M. moricandioides*; c-h, hybrids 1-6. Arrow, chloroplasts. Bar, 100 μm.

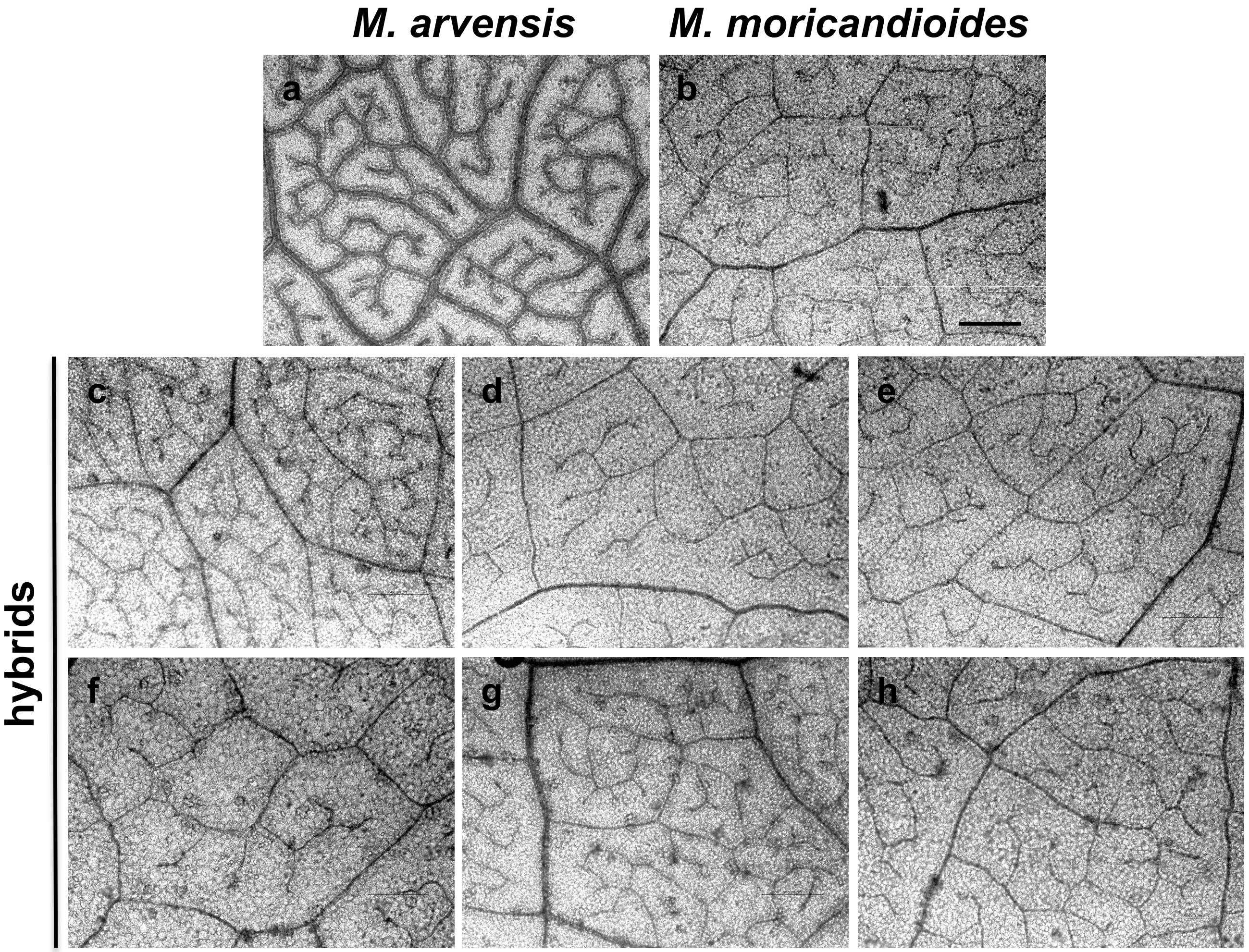


**Supplementary file 2.** Leaf venations of *M. arvensis*, *M. moricandioides* and their interspecific hybrids. a, *M. arvensis*; b, *M. moricandioides*; c-h, hybrid1-6. Bar, 500 μm.

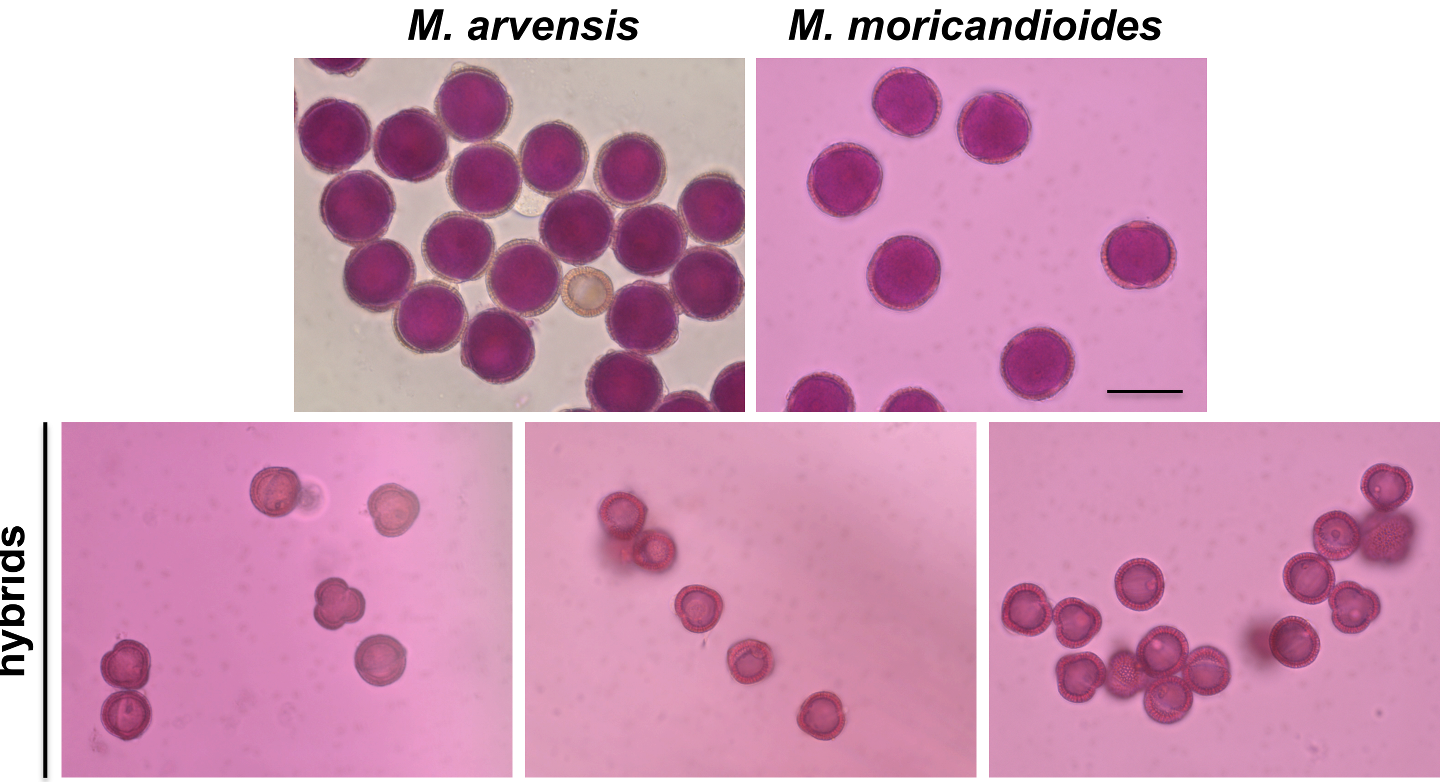


**Supplementary file 3.** The pollen activity test of *M. arvensis*, *M. moricandioides* and their interspecific hybrids dyed by Alexander staining method (Alexander, 1969). Pollens from hybrid lines were stained red, but demonstrated abnormal shapes compared to parents' round pollens. Aborted pollen grains are stained blue-green, and non-aborted pollen grains are stained magenta-red. Bar, 20 μm.

A

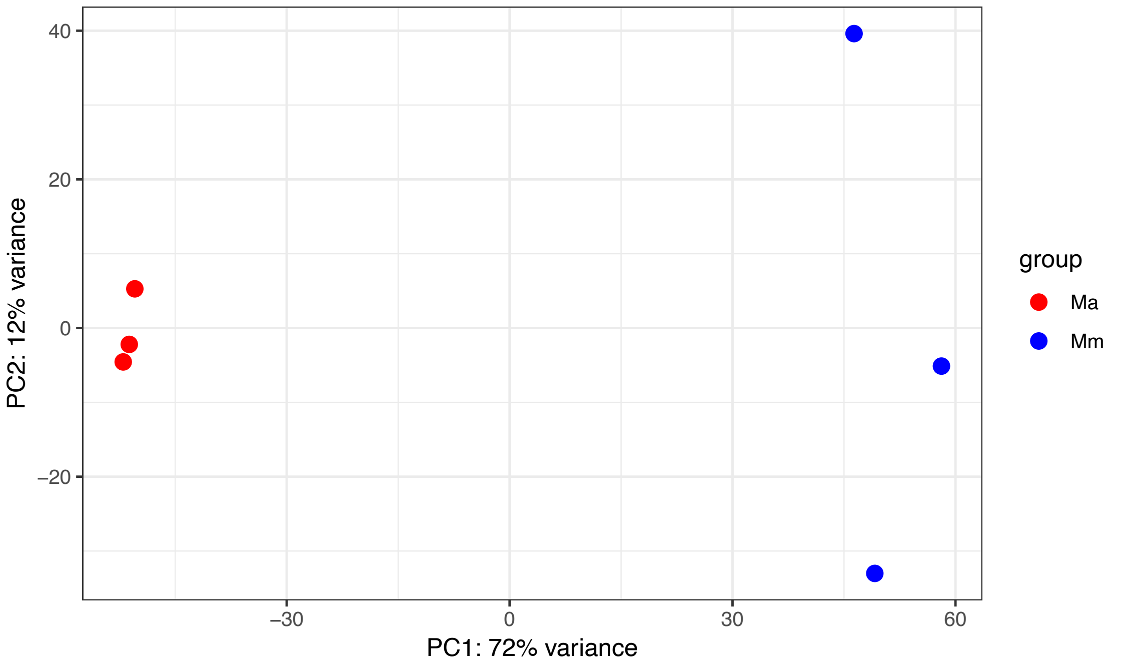


B

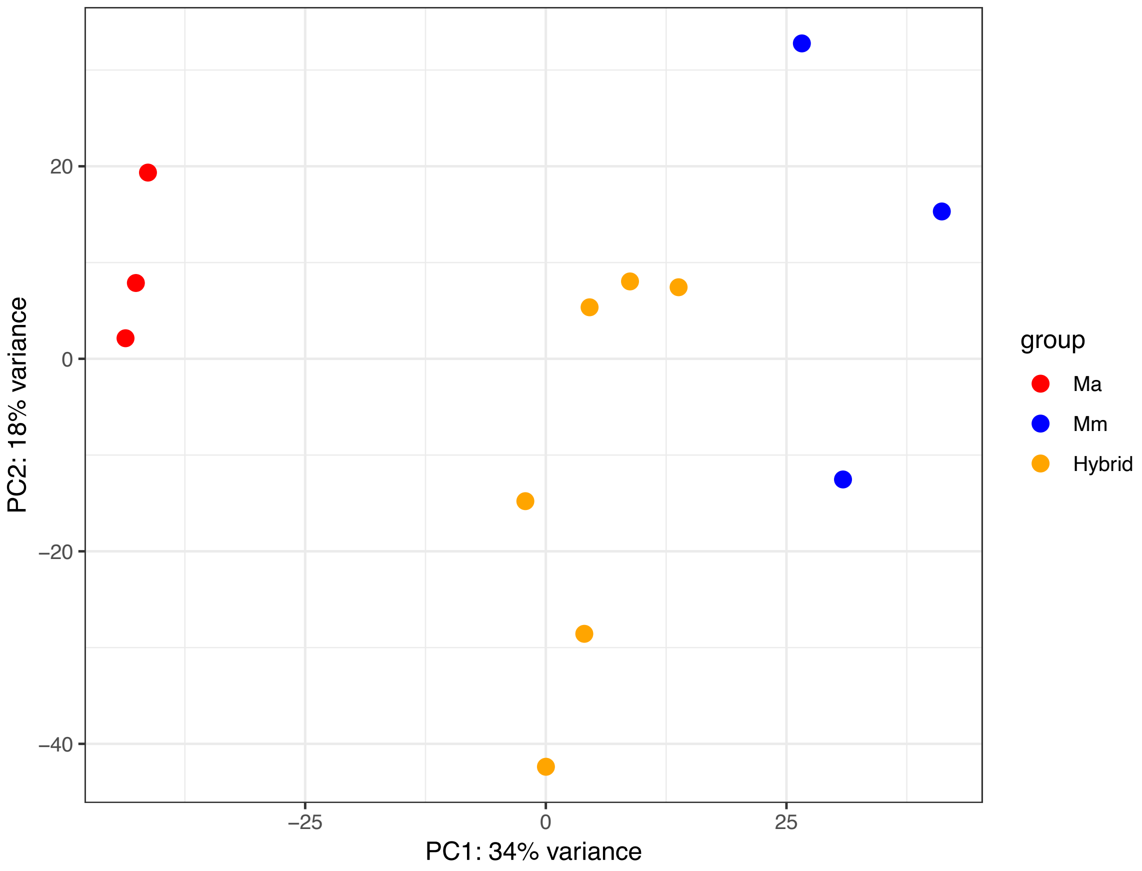


**Supplementary file 4.** Principal component analysis of rlog-transformed gene expression data A) parental species, *M. arvensis* (Ma) and *M. moricandioides* (Mm) and B) parental species and their interspecific hybrids.

**Supplementary file 5.** GO analysis on Ma-upregulated and Ma-downregulated transcripts using topGO. Top 30 most abundant GO terms were shown.

Supplementary file 5.xlsx

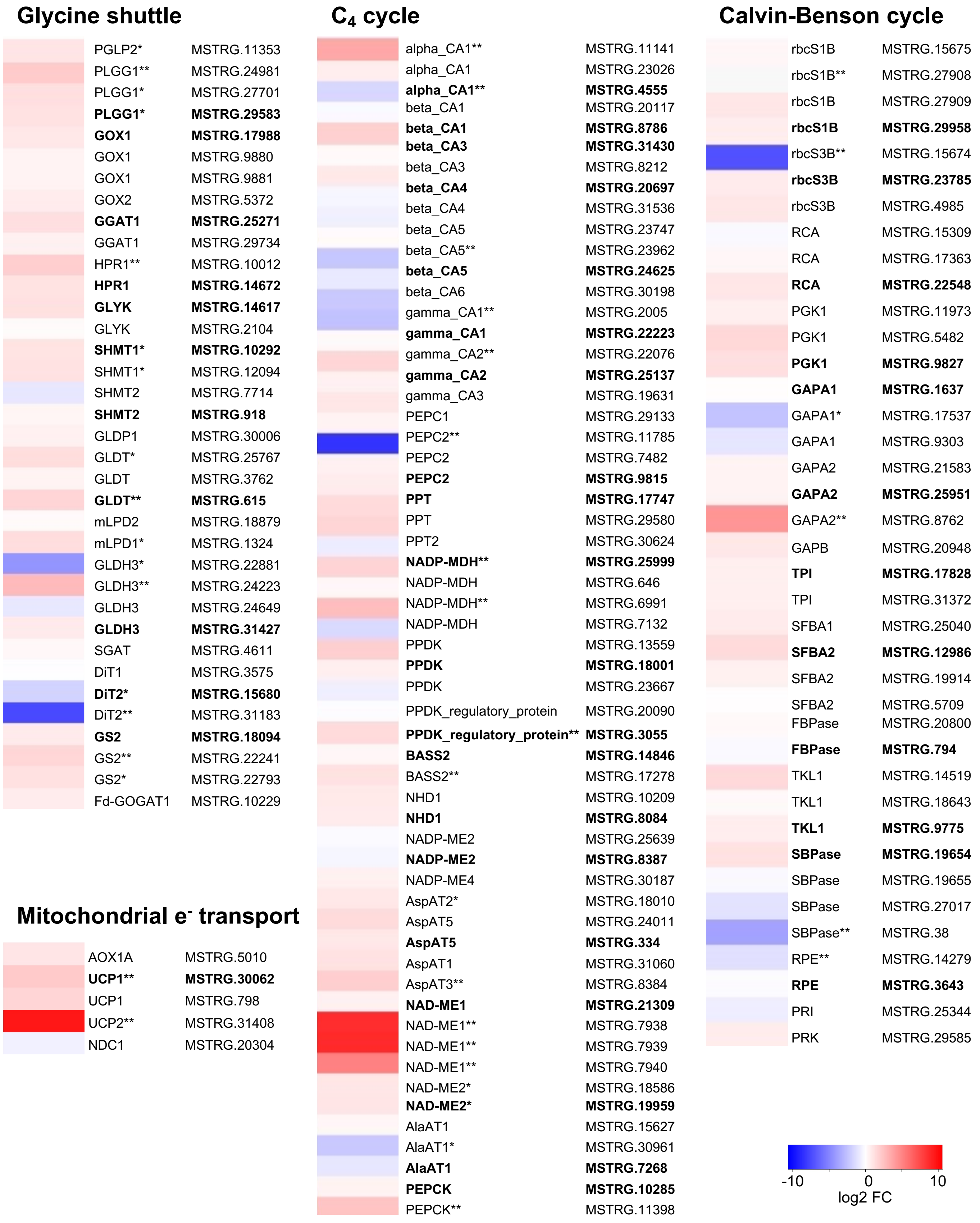


**Supplementary file 6.** Transcriptional changes in selected pathways. The heatmap indicated the log2-fold changes in transcript level of $C_3$–$C_4$ species *M. arvensis* compared to the $C_3$ species *M. moricandioides*. Blue and red indicates reduced and enhanced transcript abundance in $C_3$–$C_4$, respectively. *, adjusted P-value < 0.05; **, adjusted P-value < 0.01. Bold, transcripts with the highest expression among isoforms.

**Supplementary file 7** List of common SNPs and the transcripts harboring common SNP.

Supplementary file 7.xlsx

**Supplementary file 8.** GO analysis on common *cis*-SNPs and common *trans*-SNPs using topGO.

Supplementary file 8.xlsx

**Supplementary file 9.** Significantly enriched pathways identified in transcripts with common *cis*-SNPs and common *trans*-SNPs using KOBAS database

Supplementary file 9.xlsx

**Supplementary file 10.** Number of biased transcript with *cis*-specificity among six hybrids

| Sample name | Transcripts with *cis* -specificity | |
|---|---|---|
| | Ma_biased | Mm_biased |
| hybrid 1 | 1.105 | 820 |
| hybrid 2 | 1.128 | 822 |
| hybrid 3 | 1.050 | 758 |
| hybrid 4 | 1.086 | 793 |
| hybrid 5 | 1.124 | 847 |
| hybrid 6 | 1.057 | 825 |

**Supplementary file 11.** List of transcripts showed extreme allelic imbalance with major allele frequency ≥ 0.9 in all hybrids

| Transcript | Gene model | Bias | Description |
|---|---|---|---|
| *MSTRG.5503* | *AT5G56150* | Ma | ubiquitin-conjugating enzyme 30 |
| *MSTRG.7102* | *AT5G25120* | Ma | cytochrome p450 2C family 71 2C subfamily B 2C polypeptide 11 |
| *MSTRG.7892* | *ATCG00480* | Ma | ATP synthase subunit beta |
| *MSTRG.16149* | *AT3G14130* | Ma | Aldolase-type TIM barrel family protein |
| *MSTRG.23585* | *ATCG00490* | Ma | ribulose-bisphosphate carboxylase |
| *MSTRG.24356* | *AT4G04040* | Ma | Phosphofructokinase family protein |
| *MSTRG.27950* | *AT3G15840* | Ma | post-illumination chlorophyll fluorescence increase |
| *MSTRG.29690* | *ATCG00580* | Ma | photosystem II reaction center protein E |
| *MSTRG.29822* | *AT4G11260* | Ma | phosphatase-like protein |
| *MSTRG.5761* | *AT3G50820* | Mm | photosystem II subunit O-2 |
| *MSTRG.6356* | *AT1G13930* | Mm | oleosin-B3-like protein |
| *MSTRG.6436* | *AT5G23450* | Mm | long-chain base (LCB) kinase 1 |
| *MSTRG.6969* | *AT3G27830* | Mm | ribosomal protein L12-A |
| *MSTRG.7569* | *AT1G05010* | Mm | ethylene-forming enzyme |
| *MSTRG.7679* | *AT3G08940* | Mm | light harvesting complex photosystem II |
| *MSTRG.11356* | *AT5G47570* | Mm | NADH dehydrogenase ubiquinone 1 beta subcomplex subunit |
| *MSTRG.11905* | *AT4G05320* | Mm | polyubiquitin 10 |
| *MSTRG.15579* | *AT1G06680* | Mm | photosystem II subunit P-1 |
| *MSTRG.15675* | *AT5G38430* | Mm | Ribulose bisphosphate carboxylase (small chain) family protein |
| *MSTRG.16431* | *AT1G07970* | Mm | cytochrome B561%2C amino-terminal protein |
| *MSTRG.19717* | *AT2G27030* | Mm | calmodulin 5 |
| *MSTRG.20554* | *AT5G25280* | Mm | serine-rich protein-like protein |
| *MSTRG.20642* | *AT1G55670* | Mm | photosystem I subunit G |
| *MSTRG.23530* | *AT4G09650* | Mm | F-type H+-transporting ATPase subunit delta |
| *MSTRG.23836* | *AT1G20696* | Mm | high mobility group B3 |
| *MSTRG.24436* | *AT4G05320* | Mm | polyubiquitin 10 |
| *MSTRG.25423* | *AT1G30380* | Mm | photosystem I subunit K |
| *MSTRG.25756* | *AT5G10140* | Mm | K-box region and MADS-box transcription factor family protein |
| *MSTRG.26756* | *AT3G63410* | Mm | S-adenosyl-L-methionine-dependent methyltransferases superfamily protein |
| *MSTRG.31427* | *AT1G32470* | Mm | Single hybrid motif superfamily protein |

**Supplementary file 12.** Enrichment of regulatory effects in selected pathways. 0, no *cis*-SNP; 1, at least one *cis*-SNP found in hybrid line; 2, common *cis*-SNP among hybrids.

| Pathway | Gene | Gene name | A. thaliana | M. moricandioides | Info |
|---|---|---|---|---|---|
| **Glycine shuttle** | 2-phosphoglycolate phosphatase | PGLP2 | *AT5G47760* | *MSTRG.11353* | 1 |
| | Plastidial glycolate/glycerate transporter 1 | PLGG1 | *AT1G32080* | *MSTRG.24981* | 1 |
| | | PLGG1 | *AT1G32080* | *MSTRG.27701* | 2 |
| | | PLGG1 | *AT1G32080* | *MSTRG.29583* | 2 |
| | Glycolate oxidase | GOX1 | *AT3G14420* | *MSTRG.17988* | 1 |
| | | GOX1 | *AT3G14420* | *MSTRG.9880* | 1 |
| | | GOX2 | *AT3G14415* | *MSTRG.5372* | 1 |
| | Glutamate:glyoxylate aminotransferase | GGAT1 | *AT1G23310* | *MSTRG.25271* | 1 |
| | | GGAT1 | *AT1G23310* | *MSTRG.29734* | 2 |
| | Hydroxypyruvate reductase | HPR1 | *AT1G68010* | *MSTRG.10012* | 1 |
| | | HPR1 | *AT1G68010* | *MSTRG.14672* | 1 |
| | Glycerate kinase | GLYK | *AT1G80380* | *MSTRG.14617* | 1 |
| | | GLYK | *AT1G80380* | *MSTRG.2104* | 1 |
| | Serine hydroxymethyltransferase | SHMT1 | *AT4G37930* | *MSTRG.10292* | 1 |
| | | SHMT1 | *AT4G37930* | *MSTRG.12094* | 1 |
| | | SHMT2 | *AT5G26780* | *MSTRG.7714* | 1 |
| | | SHMT2 | *AT5G26780* | *MSTRG.918* | 1 |
| | GDC complex | GLDP1 | *AT4G33010* | *MSTRG.30006* | 2 |
| | | GLDT | *AT1G11860* | *MSTRG.25767* | 2 |
| | | GLDT | *AT1G11860* | *MSTRG.3762* | 0 |
| | | mLPD2 | *AT3G17240* | *MSTRG.18879* | 1 |
| | | mLPD1 | *AT1G48030* | *MSTRG.1324* | 1 |
| | | GLDH3 | *AT1G32470* | *MSTRG.24223* | 0 |
| | | GLDH3 | *AT1G32470* | *MSTRG.24649* | 1 |
| | | GLDH3 | *AT1G32470* | *MSTRG.31427* | 1 |
| | Serine:glyoxylate aminotransferase | SGAT | *AT2G13360* | *MSTRG.4611* | 1 |
| | Dicarboxylate transporter | DiT1 | *AT5G12860* | *MSTRG.3575* | 2 |
| | | DiT2 | *AT5G64280* | *MSTRG.15680* | 1 |
| | Plastidial glutamine synthetase | GS2 | *AT5G35630* | *MSTRG.18094* | 1 |
| | | GS2 | *AT5G35630* | *MSTRG.22241* | 0 |
| | | GS2 | *AT5G35630* | *MSTRG.22793* | 1 |
| | Ferredoxin-dependent glutamate synthase | Fd-GOGAT1 | *AT5G04140* | *MSTRG.10229* | 1 |
| **C4 cycle** | Carbonic anhydrase | alpha_CA1 | *AT3G52720* | *MSTRG.11141* | 1 |
| | | alpha_CA1 | *AT3G52720* | *MSTRG.23026* | 1 |
| | | alpha_CA1 | *AT3G52720* | *MSTRG.4555* | 2 |
| | | beta_CA1 | *AT3G01500* | *MSTRG.20117* | 1 |
| | | beta_CA1 | *AT3G01500* | *MSTRG.8786* | 1 |
| | | beta_CA4 | *AT1G70410* | *MSTRG.20697* | 2 |
| | | beta_CA4 | *AT1G70410* | *MSTRG.31536* | 1 |
| | | beta_CA5 | *AT4G33580* | *MSTRG.23747* | 2 |
| | | beta_CA5 | *AT4G33580* | *MSTRG.24625* | 1 |
| | | gamma_CA1 | *AT1G19580* | *MSTRG.22223* | 1 |
| | | gamma_CA2 | *AT1G47260* | *MSTRG.22076* | 2 |
| | | gamma_CA2 | *AT1G47260* | *MSTRG.25137* | 1 |
| | | gamma_CA3 | *AT5G66510* | *MSTRG.19631* | 1 |
| | Phosphoenolpyruvate carboxylase | PEPC1 | *AT1G53310* | *MSTRG.29133* | 1 |
| | | PEPC2 | *AT2G42600* | *MSTRG.7482* | 1 |
| | | PEPC2 | *AT2G42600* | *MSTRG.9815* | 1 |
| | Phosphoenolpyruvate/phosphate translocator | PPT | *AT5G33320* | *MSTRG.17747* | 2 |
| | | PPT2 | *AT3G01550* | *MSTRG.30624* | 1 |
| | NADP-Malate dehydrogenase | NADP-MDH | *AT5G58330* | *MSTRG.25999* | 1 |
| | | NADP-MDH | *AT5G58330* | *MSTRG.646* | 1 |
| | Pyruvate, orthophosphate dikinase | PPDK | *AT4G15530* | *MSTRG.13559* | 1 |
| | | PPDK | *AT4G15530* | *MSTRG.18001* | 1 |
| | | PPDK_regulatory_protein | *AT3G01200* | *MSTRG.20090* | 1 |
| | | PPDK_regulatory_protein | *AT4G21210* | *MSTRG.3055* | 1 |
| | | BASS2 | *AT2G26900* | *MSTRG.14846* | 2 |
| | | BASS2 | *AT2G26900* | *MSTRG.17278* | 1 |
| | | NHD1 | *AT3G19490* | *MSTRG.10209* | 2 |
| | | NHD1 | *AT3G19490* | *MSTRG.8084* | 2 |
| | NADP-malic enzyme | NADP-ME2 | *AT5G11670* | *MSTRG.25639* | 1 |
| | | NADP-ME2 | *AT5G11670* | *MSTRG.8387* | 1 |
| | | NADP-ME4 | *AT1G79750* | *MSTRG.30187* | 2 |
| | Aspartate aminotransferase | AspAT2 | *AT5G19550* | *MSTRG.18010* | 2 |
| | | AspAT5 | *AT4G31990* | *MSTRG.24011* | 1 |
| | | AspAT5 | *AT4G31990* | *MSTRG.334* | 2 |
| | | AspAT1 | *AT2G30970* | *MSTRG.31060* | 1 |
| | | AspAT3 | *AT5G11520* | *MSTRG.8384* | 2 |
| | Plasma membrane pyruvate transport | plastidic_NAD-MDH | *AT3G47520* | *MSTRG.13946* | 1 |
| | NAD-malic enzyme | NAD-ME1 | *AT2G13560* | *MSTRG.21309* | 1 |
| | | NAD-ME2 | *AT4G00570* | *MSTRG.18586* | 1 |
| | | NAD-ME2 | *AT4G00570* | *MSTRG.19959* | 1 |
| | Alanine aminotransferase | AlaAT1 | *AT1G17290* | *MSTRG.7268* | 1 |
| | Phosphoenolpyruvate carboxykinase | PEPCK | *AT4G37870* | *MSTRG.10285* | 1 |
| **Calvin-Benson cycle** | Rubisco | rbcS1B | *AT5G38430* | *MSTRG.15675* | 1 |
| | | rbcS1B | *AT5G38430* | *MSTRG.27909* | 1 |
| | | rbcS1B | *AT5G38430* | *MSTRG.29958* | 1 |
| | | rbcS3B | *AT5G38410* | *MSTRG.23785* | 1 |
| | | rbcS3B | *AT5G38410* | *MSTRG.4985* | 2 |
| | Rubisco activase | RCA | *AT2G39730* | *MSTRG.22548* | 1 |
| | Phosphoglycerate kinase | PGK1 | *AT3G12780* | *MSTRG.11973* | 1 |
| | | PGK1 | *AT3G12780* | *MSTRG.5482* | 1 |
| | | PGK1 | *AT3G12780* | *MSTRG.9827* | 1 |
| | Glyceraldehyde 3-phosphate dehydrogenase A subunit | GAPA1 | *AT3G26650* | *MSTRG.1637* | 1 |
| | | GAPA1 | *AT3G26650* | *MSTRG.9303* | 1 |
| | Glyceraldehyde 3-phosphate dehydrogenase B subunit | GAPB | *AT1G42970* | *MSTRG.20948* | 2 |
| | Triosephosphate isomerase | TPI | *AT2G21170* | *MSTRG.17828* | 2 |
| | | TPI | *AT2G21170* | *MSTRG.31372* | 1 |
| | Sedoheptulose/Fructose-biphosphate aldolase | SFBA1 | *AT2G21330* | *MSTRG.25040* | 1 |
| | | SFBA2 | *AT4G38970* | *MSTRG.12986* | 1 |
| | | SFBA2 | *AT4G38970* | *MSTRG.19914* | 1 |
| | Fructose biphosphatase | FBPase | *AT3G54050* | *MSTRG.20800* | 2 |
| | | FBPase | *AT3G54050* | *MSTRG.794* | 1 |
| | Transketolase | TKL1 | *AT3G60750* | *MSTRG.14519* | 1 |
| | | TKL1 | *AT3G60750* | *MSTRG.18643* | 1 |
| | | TKL1 | *AT3G60750* | *MSTRG.18644* | 1 |
| | | TKL1 | *AT3G60750* | *MSTRG.9775* | 2 |
| | Sedoheptulose-1,7-biphosphatase | SBPase | *AT3G55800* | *MSTRG.27017* | 0 |
| | Ribulose-5-phosphate 3-epimerase | RPE | *AT5G61410* | *MSTRG.14279* | 1 |
| | | RPE | *AT5G61410* | *MSTRG.3643* | 1 |
| | Ribulose-5-phosphate isomerase | PRI | *AT3G04790* | *MSTRG.25344* | 1 |
| | Phosphoribulokinase | PRK | *AT1G32060* | *MSTRG.24982* | 2 |
| | | PRK | *AT1G32060* | *MSTRG.27702* | 1 |
| | | PRK | *AT1G32060* | *MSTRG.29585* | 1 |
| **Mitochondrial e- transport** | Alternative oxidase | AOX1A | *AT3G22370* | *MSTRG.5010* | 1 |
| | Uncoupling protein | UCP1 | *AT3G54110* | *MSTRG.30062* | 1 |
| | | UCP1 | *AT3G54110* | *MSTRG.798* | 2 |
| | NADH dehydrogenase C1 | NDC1 | *AT5G08740* | *MSTRG.20304* | 1 |

**Supplementary file 13.** Selected gene list for promoter-GUS assay.

| Gene | Promoter region | (+1) position | Primers for promoter region amplification | | size (bp) | Final vector | Construct resistance | Agrobacteria strain | Agrobacteria resistance | Plant resistance | Primers for colony PCR | |
|---|---|---|---|---|---|---|---|---|---|---|---|---|
| *MmGLDP1* | (-)2011..(+)26 | ATG | MmGLDP1_Gibson_F | GLDP1_univ_Gibson_R | 2037 | pC1381 | Kan | GV3101(pMP90RK) | Rif, Gen | Hyg | pC1381_F | pC1381_R |
| | | | | | | | | | | | MmGLDP1_F | pC1381_R |
| *MaGLDP1* | (-)2550..(+)26 | ATG | MaGLDP1_Gibson_F | GLDP1_univ_Gibson_R | 2576 | pC1381 | Kan | GV3101(pMP90RK) | Rif, Gen | Hyg | pC1381_F | pC1381_R |
| | | | | | | | | | | | MaGLDP1_F | pC1381_R |
| *MmPHOT2* | (-)2193..(+)24 | Transcription start site | MmPHOT2_Gateway_F | MmPHOT2_Gateway_R | 2223 | pGWB3 | Kan | GV3101(pMP90RK) | Rif, Gen | Hyg, Kan | p207_F | p207_R |
| | | | | | | | | | | | pGWB3_F | pGWB3_R |
| | | | | | | | | | | | pGWB3_F | MmPHOT2_R |
| *MaPHOT2* | (-)2230..(+)3 | Transcription start site | MaPHOT2_Gateway_F | MaPHOT2_Gateway_R | 2231 | pGWB3 | Kan | GV3101(pMP90RK) | Rif, Gen | Hyg, Kan | p207_F | p207_R |
| | | | | | | | | | | | pGWB3_F | pGWB3_R |
| | | | | | | | | | | | pGWB3_F | MaPHOT2_R |
| *MmCHUP1* | (-)2000..(-)1 | Putative ATG | MmCHUP1_Gibson_F | MmCHUP1_Gibson_R | 2000 | pC1381 | Kan | GV3101(pMP90RK) | Rif, Gen | Hyg | pC1381_F | pC1381_R |
| | | | | | | | | | | | MmCHUP1_F | pC1381_R |
| *MaCHUP1* | (-)2000..(-)1 | Putative ATG | MaCHUP1_Gibson_F | MaCHUP1_Gibson_R | 2000 | pC1381 | Kan | GV3101(pMP90RK) | Rif, Gen | Hyg | pC1381_F | pC1381_R |
| | | | | | | | | | | | MaCHUP1_F | pC1381_R |
| *MmDUF538* | (-)2491..(+)23 | Putative ATG | MmDUF538_Gibson_F | DUF538_univ_Gibson_R | 2514 | pC1381 | Kan | GV3101(pMP90RK) | Rif, Gen | Hyg | pC1381_new_F | pC1381_new_R |
| | | | | | | | | | | | MmDUF538_F | pC1381_new_R |
| *MaDUF538* | (-)2231..(+)23 | Putative ATG | MaDUF538_Gibson_F | DUF538_univ_Gibson_R | 2254 | pC1381 | Kan | GV3101(pMP90RK) | Rif, Gen | Hyg | pC1381_new_F | pC1381_new_R |
| | | | | | | | | | | | MaDUF538_F | pC1381_new_R |
| *MmATPB* | (-)2301..(+)28 | Putative ATG | MmATPB_Gibson_F | ATPB_univ_Gibson_R | 2329 | pC1381 | Kan | GV3101(pMP90RK) | Rif, Gen | Hyg | pC1381_new_F | pC1381_new_R |
| | | | | | | | | | | | ATPB_F | pC1381_new_R |
| *MaATPB* | (-)2293..(+)28 | Putative ATG | MaATPB_Gibson_F | ATPB_univ_Gibson_R | 2321 | pC1381 | Kan | GV3101(pMP90RK) | Rif, Gen | Hyg | pC1381_new_F | pC1381_new_R |
| | | | | | | | | | | | ATPB_F | pC1381_new_R |

**Supplementary file 14.** Confirmation of RNA-Seq data by allele-specific RT-PCR of *M. arvensis* × *M. moricandioides* hybrid.

| *Moricandia* gene | Hybrid | Allele-specific transcript level and allelic ratio in RNA-Seq | | | Allelic ratio in qPCR |
|---|---|---|---|---|---|
| | | Ma allele in F1 | Mm allele in F1 | Allelic ratio Ma:Mm in F1 | Allelic ratio Ma:Mm in F1 |
| *GLK2* | 1 | 77 | 145 | 0.53 | 0.18 |
| (*MSTRG.28909*) | 2 | 52 | 151 | 0.34 | 0.37 |
| | 3 | 38 | 128 | 0.30 | 0.21 |
| | 4 | 67 | 122 | 0.55 | 0.12 |
| | 5 | 61 | 148 | 0.41 | 0.18 |
| | 6 | 87 | 137 | 0.64 | 0.20 |
| *ASP3* | 1 | 283 | 74 | 3.82 | 35.39 |
| (*MSTRG.8384*) | 2 | 404 | 100 | 4.04 | 58.21 |
| | 3 | 180 | 65 | 2.77 | 31.86 |
| | 4 | 466 | 88 | 5.30 | 1.30 |
| | 5 | 295 | 51 | 5.78 | 41.20 |
| | 6 | 395 | 70 | 5.64 | 55.84 |
| γCA2 | 1 | 180 | 85 | 2.12 | 3.10 |
| (*MSTRG.22076*) | 2 | 173 | 89 | 1.94 | 5.20 |
| | 3 | 121 | 47 | 2.57 | 3.01 |
| | 4 | 151 | 46 | 3.28 | 3.82 |
| | 5 | 159 | 66 | 2.41 | 3.74 |
| | 6 | 113 | 61 | 1.85 | 2.37 |
| *PPA2* | 1 | 299 | 121 | 2.47 | 2.57 |
| (*MSTRG.29297*) | 2 | 331 | 170 | 1.95 | 5.72 |
| | 3 | 171 | 85 | 2.01 | 2.83 |
| | 4 | 713 | 359 | 1.99 | 2.41 |
| | 5 | 380 | 173 | 2.20 | 2.24 |
| | 6 | 346 | 158 | 2.19 | 2.58 |

**Supplementary file 15.** qPCR primer list for ASE verification.

| Gene | Ortholog *A. thalinana* | Ortholog *M. moricandioides* | Primer name | Sequence (5'-3') | Product size (bp) |
|---|---|---|---|---|---|
| *Helicase* | *AT1G58060* | *MSTRG.22364* | mori_Helicase_10F | CGGATGCCATTGGTAGAACT | 97 |
| | | | mori_Helicase_10R | CTTCACTCGGAGGTTCCAAA | |
| *GLK2* | *AT5G44190* | *MSTRG.28909* | GLK2_SNP1_a_2R | CGATCATCTTCACCGTCATAG | 247 |
| | | | GLK2_SNP1_m_2R | CGATCATCTTCACCGTCATAA | |
| | | | GLK2_SNP1_g_2F | ATCGGAAACGTCAAAGGGAT | |
| *ASP3* | *AT5G11520* | *MSTRG.8384* | ASP3_SNP2_a_1F | AATGTACTCAAATCCTCCGAGC | 152 |
| | | | ASP3_SNP2_m_1F | AATGTACTCAAATCCTCCGAGT | |
| | | | ASP3_SNP2_g_1R | GAGCACGTAATGCCTCGAA | |
| *γCA2* | *AT1G47260* | *MSTRG.22067* | gamma_CA2_SNP2_a_1F | CTTCTCAACCACCACACCATAA | 57 |
| | | | gamma_CA2_SNP2_m_1F | CTTCTCAACCACCACACCATAG | |
| | | | gamma_CA2_SNP2_g_1R | GATGAGGCATTTGTTGGCAT | |
| | | | gamma_CA2_SNP2_a_2F | TTCTCAACCACCACACCATAA | 63 |
| | | | gamma_CA2_SNP2_m_2F | TTCTCAACCACCACACCATAG | |
| | | | gamma_CA2_SNP2_g_2,3R | TGTTGAGGATGAGGCATTTG | |
| | | | gamma_CA2_SNP2_a_3F | TCTCAACCACCACACCATAA | 62 |
| | | | gamma_CA2_SNP2_m_3F | TCTCAACCACCACACCATAG | |
| *PPA2* | *AT2G18230* | *MSTRG.29297* | PPA2_SNP2_a_1F | CATTCCTCGAACCATTTGTGTA | 194 |
| | | | PPA2_SNP2_m_1F | CATTCCTCGAACCATTTGTGTG | |
| | | | PPA2_SNP2_g_1R | GAAGGAATGACCCAGTCAGC | |

**Supplementary file 16.** Primer list for promoter-GUS assay.

| Primer name | Sequence (5'-3') |
|---|---|
| MmGLDP1_Gibson_F | CGGCGCGCCGAATTCCCGGGGATCCCCGGTAACTCTTTAAATTGCTTG |
| MaGLDP1_Gibson_F | CGGCGCGCCGAATTCCCGGGGATCCGGAGCGGAACTCTTACGAG |
| GLDP1_univ_Gibson_R | CGTAAACTAGTCAGATCTACCATGGTAAGCAAGCCTACGTGCG |
| MmPHOT2_Gateway_F | GGGGACAAGTTTGTACAAAAAAGCAGGCTTCGCACTATCATTCCTCACCAT |
| MmPHOT2_Gateway_R | GGGGACCACTTTGTACAAGAAAGCTGGGTCGAAGGACCACACACTCTGTT |
| MaPHOT2_Gateway_F | GGGGACAAGTTTGTACAAAAAAGCAGGCTTCGACAAAGGCAGAAGACTGAC |
| MaPHOT2_Gateway_R | GGGGACCACTTTGTACAAGAAAGCTGGGTCTCCTTTCCTCTTTTTACTCC |
| MmCHUP1_Gibson_F | CGGCGCGCCGAATTCCCGGGGATCCTAGAATCTCTGCTTTGATAAAAG |
| MmCHUP1_Gibson_R | CGTAAACTAGTCAGATCTACCATGGATATTAACACCTTGAATTGTGAATAAC |
| MaCHUP1_Gibson_F | CGGCGCGCCGAATTCCCGGGGATCCAGAGGCTAACAACGGATAAATC |
| MaCHUP1_Gibson_R | CGTAAACTAGTCAGATCTACCATGGATATTAACACCTTGAATTATGAATAAAC |
| MmDUF538_Gibson_F | CGGCGCGCCGAATTCCCGGGGATCCGCGATATTGGGCTTTTGTG |
| MaDUF538_Gibson_F | CGGCGCGCCGAATTCCCGGGGATCCCATGGCGTTGCTTATGGG |
| DUF538_univ_Gibson_R | CGTAAACTAGTCAGATCTACCATGGTGCATCATCATCATCTTCG |
| MmATPB_Gibson_F | CGGCGCGCCGAATTCCCGGGGATCCGTTCCTGTTCAGCTTCTTGAG |
| MaATPB_Gibson_F | CGGCGCGCCGAATTCCCGGGGATCCGTTCATGTTTGTGGATCACAAG |
| ATPB_univ_Gibson_R | CGTAAACTAGTCAGATCTACCATGGCCGCCACTTTCTTACCCG |
| p207_F | TCGCGTTAACGCTAGCATGGATCTC |
| p207_R | GTAACATCAGAGATTTTGAGACAC |
| pGWB3_F | GCCTGCAGGTCGACTCTAAT |
| pGWB3_R | GGTTGGGGTTTCTACAGGAC |
| pC1381_F | CGTGCTCCACCATGTTGG |
| pC1381_R | CTGCATCGGCGAACTGATC |
| pC1381_new_F | CCACCATGTTATCACATCAA |
| pC1381_new_R | CCCGCATAATTACGAATATC |
| MmGLDP1_F | CATTTTCGTCCACCAAATCC |
| MaGLDP1_F | CCCAGCTCGCTTCTCAAGTA |
| MmPHOT2_R | CTCCTCAGGAAGCTCATGCT |
| MaPHOT2_R | TCTTGGAGTTGGGACTTCGT |
| MmCHUP1_F | ATTTCACGAACTGGGTTTGC |
| MaCHUP1_F | CCACTTCCTCCTCCTCCTCT |
| MmDUF538_F | CCACTAGGGTCATGTTCATT |
| MaDUF538_F | CAAGGACTGATGCATACAAA |
| ATPB_F | CTTGCTTCTCCTCTTCCTCT |
